# Identification and characterization of a bacterial periplasmic solute binding protein that binds L-amino acid amides

**DOI:** 10.1101/2024.02.22.581678

**Authors:** Oliver B. Smith, Rebecca L. Frkic, Marina G. Rahman, Colin J. Jackson, Joe A. Kaczmarski

**Affiliations:** Research School of Chemistry, Australian National University, Canberra, ACT 2601, Australia; ARC Centre of Excellence in Synthetic Biology, Australian National University, Canberra, ACT 2601, Australia; ARC Centre of Excellence for Innovations in Peptide & Protein Science, Australian National University, Canberra, ACT 2601, Australia; Research School of Biology, Australian National University, Canberra, ACT 2601, Australia

**Author notes:** Corresponding author(s): Joe Kaczmarski,; Colin Jackson.

## Abstract

Periplasmic solute-binding proteins (SBPs) are key ligand recognition components of bacterial ATP-binding cassette (ABC) transporters that allow bacteria to import nutrients and metabolic precursors from the environment. Periplasmic SBPs comprise a large and diverse family of proteins, of which only a small number have been empirically characterized. In this work, we identify a set of 610 unique uncharacterized proteins within the SBP_bac_5 family that are found in conserved operons comprising genes encoding (i) ABC transport systems and (ii) putative amidases from the FmdA_AmdA family. From these uncharacterized SBP_bac_5 proteins, we characterize a representative periplasmic SBP from *Mesorhizobium* sp. A09 (*Me*Ami_SBP) and show that *Me*Ami_SBP binds l-amino acid amides but not the corresponding l-amino acids. An X-ray crystal structure of *Me*Ami_SBP bound to l-serinamide highlights the residues that impart distinct specificity for l-amino acid amides and reveals a structural Ca^2+^ binding site within one of the lobes of the protein. We show that the residues involved in ligand and Ca^2+^ binding are conserved amongst the 610 SBPs from experimentally uncharacterized FmdA_AmdA amidase-associated ABC transporter systems, suggesting these homologous systems are also likely to be involved in the sensing, uptake and metabolism of l-amino acid amides across many Gram-negative nitrogen-fixing soil bacteria. We propose that *Me*Ami_SBP is involved in the uptake of such solutes to supplement pathways such as the citric acid cycle and the glutamine synthetase-glutamate synthase pathway. This work expands our currently limited understanding of microbial interactions with l-amino acid amides and bacterial nitrogen utilization.

## INTRODUCTION

Bacterial ATP-binding cassette (ABC) transporter systems enable bacteria to adapt to diverse environments by facilitating the selective uptake of key nutrients and metabolic precursors (1). While bacterial ABC transporter systems for sugars (2–5), oligopeptides (6,7) and amino acids (8–10) are well characterized, those specialized for the import and utilization of other important carbon and nitrogen sources, such as amides, remain poorly understood.

In Gram-negative bacteria, ABC transporter systems are typically encoded by a single operon that include genes for (i) a periplasmic solute binding protein (SBP) that initiates solute recognition and capture, (ii) transmembrane permease and ATP-binding domains for the transport of the solute into the cytoplasm and, in some cases, (iii) cytosolic enzyme(s) that catalyze the first step in solute metabolism (1,11). Because the periplasmic SBP components determine the ligand specificity of the transport system (12–15), identifying uncharacterized SBPs with novel ligand binding specificities can help expand our understanding of bacterial ABC transporters and their role in solute uptake and metabolism.

The SBP superfamily is a large group of proteins (> 3.4 million sequences in February 2024) (16,17) with diverse sequences and ligand-binding preferences (13,18). Although prokaryotic SBPs share a common core architecture comprising two primary α/β domains linked by a flexible hinge region and a central ligand-binding cleft, their structures are diverse and cover 33 distinct Pfam structural families (19). Proteins of the “extracellular solute-binding protein, family 5” Pfam family (“SBP_bac_5”, PF00496) are amongst the most structurally divergent SBPs and contain three topological domains (13). Unlike other structural classes of SBPs, which show preferences for a particular ligand class (19), members of the SBP_bac_5 family bind a range of ligand types, including oligopeptides (20,21), glutathione (22,23) and metal ions (24,25). While a handful of the SBP_bac_5 proteins have been experimentally characterized, much of the sequence and functional diversity of the SBP_bac_5 family remains unexplored.

In this work, we use sequence similarity network (SSN) and genomic context analyses to identify a cluster of 610 unique uncharacterized proteins within the SBP_bac_5 family that are all found within ABC transporter operons that also contain an enzyme from the FmdA_AmdA acetamidase/formamidase family. We experimentally characterize a representative of this group of proteins, *Me*Ami_SBP from *Mesorhizobium* sp. AP09, and show that *Me*Ami_SBP binds l-amino acid amides but not the corresponding l-amino acids. A crystal structure of *Me*Ami_SBP bound to l-serinamide highlights the key interactions and binding site properties that confer this novel, divergent, ligand specificity and reveals a Ca^2+^ binding site in one of the protein’s lobes. Ligand-binding residues are conserved amongst the other proteins in this cluster, suggesting that this group of proteins has a role in the uptake and metabolism of l-amino acid amides. This work expands our limited understanding of microbial interactions with l-amino acid amides, especially in relation to bacterial nitrogen utilization.

## RESULTS

### Identification of a group of uncharacterized SBPs from amidase-containing ABC transporter operons in Gram-negative bacteria

Sequence similarity networks (SSNs) are valuable tools for identifying and exploring uncharacterized regions of sequence-function space within protein families (26,27). We generated an SSN comprising ∼400,000 protein sequences from the SBP_bac_5 family (**Figure 1A**). Mapping SWISS-PROT annotations and PDB structures onto this SSN show that the experimentally characterized members of the SBP_bac_5 family are concentrated within localized regions of the sequence space. Indeed, when the SSN was generated using an alignment score cut-off of 100 (corresponding to roughly 30% sequence identity) most of the distinct sequence clusters had no experimentally characterized members, highlighting that much of the functional diversity of this family remains unexplored.

**Figure 1.**
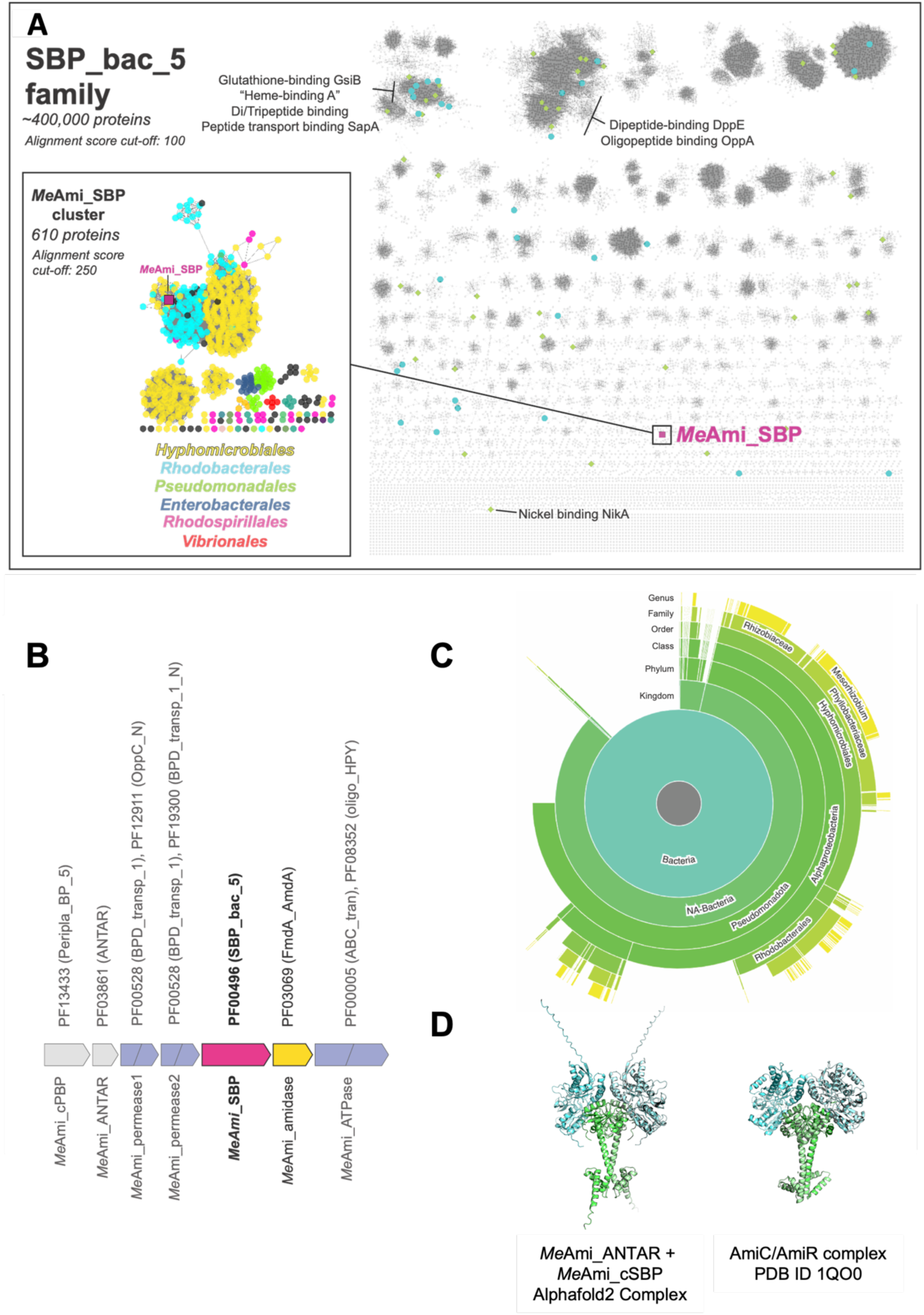
A group of 610 homologous SBP_bac_5 solute binding proteins from conserved amidase-containing ABC transporter operons. **A.** A sequence similarity network (SSN) of the SBP_bac_5 Pfam using an alignment cut-off score of 100. Nodes representing proteins with published PDB structures are shown as green diamonds. Nodes representing proteins with reviewed SWISS-PROT annotations (but no PDB structures) are shown as cyan circles. The node representing *Me*Ami_SBP is shown as a magenta square. A few key SWISS-PROT annotations are labelled. **(insert)** A close-up of the *Me*Ami_SBP-containing cluster, using an alignment cut-off score of 250. Nodes are colored based on the Order of the host organism (except for *Me*Ami_SBP, magenta square). **B.** The conserved genomic neighborhood/operon structure amongst *Me*Ami_SBP and its homologs. Each open-reading frame is labelled above with the corresponding Pfam identifiers. Each gene is also labelled with the name we give the corresponding gene in the *Me*Ami operon from *Mesorhizobium* sp. AP09. **C.** The taxonomic distribution of proteins in the *Me*Ami_SBP cluster generated using the EFI-EST Taxonomy tool (46). **D.** An AF2-predicted model of the *Me*Ami_ANTAR/*Me*Ami_cPBP complex (left) and the X-ray crystal structure of the *P. aeruginosa* AmiC/AmiR complex (right). In each structure, the respective PBP components are shown in light blue, and the respective ANTAR components are shown in green.

To identify novel and interesting ligand binding properties within the SBP_bac_5 family, we performed genomic context analyses of each SBP_bac_5 protein cluster to identify which SBP open reading frames were colocalized with ABC transporter cassettes and were also found near genes encoding enzymes that could suggest a biochemical requirement for processing novel ligands (28). One such group comprised 610 unique SBP_bac_5 proteins that shared a highly conserved ABC transporter operon structure containing a gene that encodes a protein from the acetamidase/formamidase family (PF03069, FmdA_AmdA) (**Figure 1B**).

The presence of a putative FmdA_AmdA amidase family member in each of these operons suggested that these ABC transport systems may be involved in the import and processing of amine-containing compounds. While the FmdA_AmdA family of proteins is primarily known for hydrolysis of acetate (29) and formate (30), some members of this family have activity on other amide-containing compounds, including lactams and aliphatic amides (31). Interestingly, the cluster of 610 amidase-associated operons that we identified is dominated by proteins from Gram-negative bacteria that are specialized in the uptake of nitrogen-containing compounds, such as *Rhizobiales, Mesorhizobium* and *Rhodobacteraceae* (32–34) (**Figure 1A, 1C**). Notably, no other clusters within the SBP_bac_5 family were associated with a FmdA_AmdA protein and the structure of these amidase-associated operons was clearly distinct from previously well-studied amidase operons such as the *Pseudomonas aeruginosa* amidase operon (35,36), *Rhodococcus* sp. R312 amidase operon (37) and *Mycobacterium smegmatis* acetamidase operon (38–40) (**Supplementary Figure 1**). This marked difference prompted further investigation into this distinct set of ABC transporter-associated proteins.

In addition to genes for the putative amidase and the core ABC transporter import proteins (i.e. the periplasmic SBP, membrane permease and ATP-binding domains), two additional genes were consistently present at the 5’ end of each operon in this cluster. These genes were predicted to encode (i) a protein with a periplasmic binding protein fold (PF13433, Peripla_BP_5) that lacks a periplasmic signaling peptide and (ii) a protein belonging to the family of AmiR and NasR transcription antitermination regulators (ANTAR, PF03861) (**Figure 1B**). The presence of transcription antitermination regulators in all of these operons suggested that the expression of these ABC transporter and associated proteins may be regulated in a ligand-dependent ANTAR-mediated manner like the amide import operon of *P. aeruginosa* (41). This potential regulatory mechanism also prompted investigation into the SBP component of this conserved operon.

To establish the function and substrate specificity of this group of ABC transporters associated with potentially unique enzymatic processing and regulatory mechanisms, we focused on identifying the ligand specificity of the associated periplasmic SBPs. Rather than directly experimentally characterizing each member of this cluster (which share over 35% sequence identity), we used a profile hidden Markov model (43) to identify the most consensus-like, or representative, SBP sequence within this group of proteins. The selected representative protein, which we call *Me*Ami_SBP, comes from the soil bacterium *Mesorhizobium* sp. AP09 and is annotated on UniProt as being a “peptide/nickel transport system substrate-binding protein/oligopeptide transport system substrate-binding protein” (UniProt ID: A0A395KPD0) (44). Importantly, *Me*Ami_SBP has no closely related homologs that have been experimentally characterized, with the closest related protein with a confirmed function being the glutathione-binding protein, GsiB, from *Salmonella enterica* (BLAST E-value = 9e^-73^, 32% sequence identity) (45). Of note, the ANTAR protein in the *Me*Ami operon (*Me*Ami_ANTAR) shares 32% sequence identity with the *P. aeruginosa* ANTAR protein, AmiR, and the cytosolic Peripla_BP_5 protein from the *Me*Ami operon (*Me*Ami_cPBP) shares 29% sequence identity with the cytosolic ANTAR-associated ligand receptor from *P. aeruginosa*, AmiC. An Alphafold2 (42) model of these two proteins predict they will form a dimer of dimers similar to the AmiC/AmiR complex (**Figure 1D**).

### *Me*Ami_SBP binds l-amino acid amides

His_6_-tagged *Me*Ami_SBP was expressed in the cytosol of *Escherichia coli* BL21 (DE3) cells and purified. *Me*Ami_SBP is monomeric in solution (**Supplementary Figure 2**). To identify ligands of *Me*Ami_SBP, we screened for binding to 510 biologically-relevant compounds from the Biolog Phenotype Microarray plates PM1-PM6 using a differential scanning fluorimetry (DSF)-based thermal shift assay. The PM1-PM6 screens are designed for use in bacterial phenotyping studies that assess the uptake and metabolism of common solutes (47), and therefore contain compounds that ABC transporter-associated SBPs, such as *Me*Ami_SBP, are likely to bind **(Supplementary Tables 1-6).**

Despite *Me*Ami_SBP being annotated as a peptide binding protein, no thermal shifts were observed for any of the ∼100 di- and tripeptides present in the screen (**Supplementary Figure 3**). Similarly, there were no signs of ligand-induced stabilization by other known ligands of the SBP_bac_5 family, such as glutathione and octopine. A clear ligand-induced thermal shift (ι1T_m_ 14.1 °C compared to negative control) was observed in the presence of an l-amino acid amide, L-alaninamide, only (**Figure 2A**). L-alaninamide was the only amino acid amide in the initial screen; however, subsequent experiments also showed ligand-induced thermal shifts in the presence of 10 mM glycinamide and L-tyrosinamide, indicating *Me*Ami_SBP binds l-amino acid amides with diverse side chain chemistry (**Figure 2B**). Isothermal analysis of ligand-titrated DSF experiments revealed that the affinity of *Me*Ami_SBP for the three amino acid amides spans two orders of magnitude: glycinamide (*K*_D_ of 49.7 μM), L-alaninamide (*K*_D_ of 1.53 μM) and L-tyrosinamide (*K*_D_ of 0.187 μM) (**Figure 2C, 2D**) (48).

**Figure 2.**
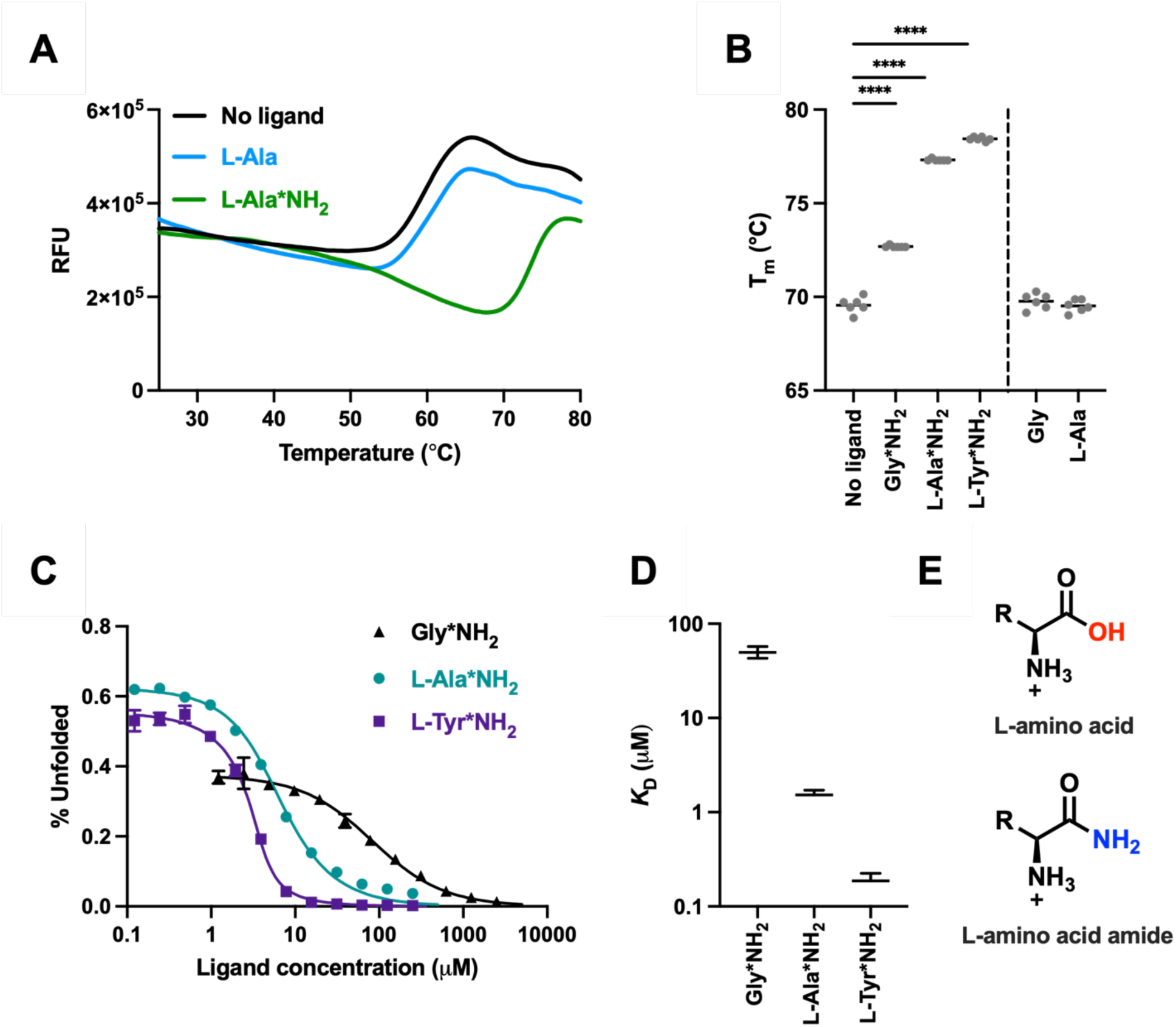
*Me*Ami_SBP from *Mesorhizobium* sp. A09 is a periplasmic solute binding protein specific for l-amino acid amides. **A.** Representative DSF fluorescence traces of *Me*Ami_SBP in the presence of L-alanine, L-alaninamide (L-Ala*NH_2_), and a no ligand control. RFU = relative fluorescent units. **B.** Melting temperatures (T_m_) of *Me*Ami_SBP in the presence of 10 mM of various amino acid amides (L-X*NH_2_) and the corresponding amino acids as measured by DSF (n=6, horizontal line represents the mean). **** refers to p < 0.0001 from unpaired parametric t-tests. **C.** Isothermal DSF curves from amino acid amide titrations into *Me*Ami_SBP. Data is (n=3, error bars=SD). A one-site model (solid line) was fitted to the data using FoldAffinity (48). **D.** Binding affinities of *Me*Ami_SBP for the l-amino acid amides. Data represent the mean, lower and upper limit of *K*_D_ values from the fit of the data in (C). **E.** General chemical structures of l-amino acids and l-amino acid amides. R= side chain (i.e. Gly: R=H, Ala: R=CH_3_, etc.).

Importantly, thermal shifts were not observed for any of the proteinogenic l-amino acids tested (**Figure 2B**), confirming selectivity for the l-amino acid amide moiety over the l-amino acid moiety (**Figure 2E**). Similarly, no thermal shifts were observed in the presence of non-amino acid amides such as acetamide and glucuronamide (**Supplementary Figure 3**). These data raised questions about (i) the structural basis of selectivity for l-amino acid amides over l-amino acids and other amidated compounds and (ii) the structural basis for large differences in ligand affinity that could be attributed solely to differences in side chain chemistry.

### The structure of *Me*Ami_SBP in complex with an amino acid amide

We obtained a 1.55 Å crystal structure of *Me*Ami_SBP, heterologously expressed and purified from *Escherichia coli*. As expected, *Me*Ami_SBP adopts the SBP_bac_5 fold, with three subdomains (**Figure 3A**, **Supplementary Table 7**).

**Figure 3.**
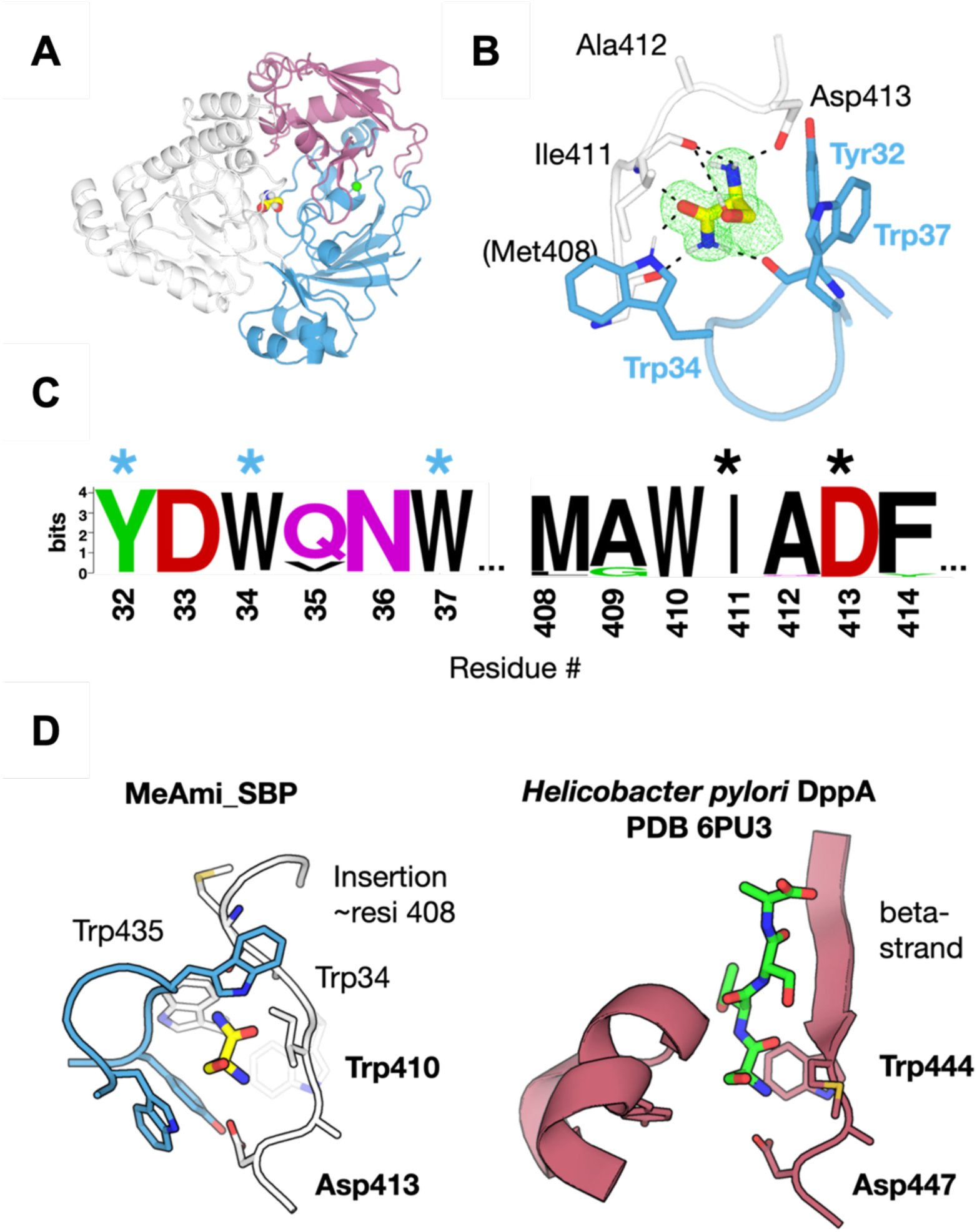
Structural analysis of *Me*Ami_SBP supports specific binding to amino acid amides. **A.** X-ray crystal structure of *Me*Ami_SBP showing the three subdomains of the protein (white, pink and blue), the bound l-serinamide molecule (spheres) within the central binding site, and the bound Ca^2+^ ion (green sphere). **B.** The amino acid amide binding site of *Me*Ami_SBP showing key interactions. Hydrogens were added in ChimeraX to emphasize the H-bonding network. A Polder omit map around the ligand is shown as a green mesh (Contoured at 3 sigma, 3 Å carve around ligand). **C.** Sequence WebLogo (52) showing sequence conservation amongst *Me*Ami_SBP homologs at residue positions important for binding amino acid amides in *Me*Ami_SBP. Residues whose sidechains interact with the l-serinamide ligand in *Me*Ami_SBP are highlighted with an asterisk, with blue asterisks referring to residues on the small core lobe of *Me*Ami_SBP, and black asterisks referring to residues on the large core lobe of *Me*Ami_SBP. **D.** Comparison of the binding cavities of *Me*Ami_SBP and the dipeptide binding protein, DppA, from *Helicobacter pylori* (PDB 6PU3), highlighting similarities (e.g. positioning of Asp413/447) and differences (e.g. disruption of beta-strand in *Me*Ami_SBP) that contribute to differing ligand specificities.

Clear electron density within the canonical SBP binding cleft indicated that *Me*Ami_SBP had co-purified and crystallized bound to a ligand from the expression host. The high resolution of the data meant that we could confidently model l-serinamide within the binding site (**Figure 3B**). Analysis of the structure of *Me*Ami_SBP in complex with l-serinamide revealed key interactions between the protein and the ligand. This includes (i) a salt-bridge between the N-terminal amine of l-serinamide and Asp413, (ii) hydrogen bonds between the l-serinamide carbonyl and both the NH group of the Trp34 sidechain and the amide NH of Ile411, and (iii) hydrogen bonds between the nitrogen of l-serinamide’s carboxamide group and the backbone carbonyl groups of Tyr32 and Met408.

The structure of this complex helps to explain the selectivity of *Me*Ami_SBP for amino acid amides over the corresponding l-amino acids: the hydrogen bonds observed between the carboxamide group of l-serinamide and *Me*Ami_SBP would be disrupted if the carboxamide group was replaced with a carboxylic acid (as would be the case for the corresponding amino acids). Similarly, the key interaction between the N-terminal amino group of l-serinamide and Asp413 explains why *Me*Ami_SBP does not bind amides that lack an N-terminal amino group, such as acetamide and glucuronamide.

Given that our DSF experiments showed that *Me*Ami_SBP binds glycinamide, L-alaninamide and L-tyrosinamide with various affinities, we used computational docking to predict and understand how these amino acid amides would bind *Me*Ami_SBP. As expected, top-scoring poses show glycinamide, L-alaninamide and L-tyrosinamide occupying the same binding site in *Me*Ami_SBP as l-serinamide, with the backbone motif held in place by the same interactions observed in the l-serinamide complex (**Supplementary Figure 4**). With the backbone of the l-amino acid amides held in this position, the side chains point away from the binding site where there is space within the binding cleft of *Me*Ami_SBP to accommodate bulkier sidechains, such as that of L-tyrosinamide. While most of the interactions between the ligands and the protein are from the l-amino acid amide backbone, additional interactions with the ligand side chains likely contribute to the observed differences in binding affinity between glycinamide, L-alaninamide and L-tyrosinamide. For example, glycinamide, which had the lowest binding affinity in isothermal DSF experiments (*K*_D_ of 49.7 μM) lacks a side chain and thus forms fewer stabilizing interactions. On the other hand, the side chain of L-tyrosinamide likely forms hydrophobic and pi-stacking interactions with Trp32, Trp410, Trp435 and Tyr32, which would contribute to the increased affinity (*K*_D_ of 0.187 μM). The lack of water molecules in the side chain-binding site (**Figure 3B**) indicates side chain contributions to l-amino acid amide binding are primarily enthalpic, with no entropic contribution being made by the expulsion of water molecules from this binding site.

Sequence analysis of the related SBP proteins from the same cluster as *Me*Ami_SBP reveals that all residues involved in binding l-amino acid amide backbone in *Me*Ami_SBP are highly conserved (**Figure 3C**), suggesting that other proteins within this cluster likely have a similar ligand binding profile to *Me*Ami_SBP. This, together with the conserved operon structure amongst all 610 proteins in this group of proteins, suggests that this distinct group of amidase-associated ABC transporters share a common substrate and physiological function.

We also compared the structure of *Me*Ami_SBP with published structures of other members of the SBP_bac_5 family, including the most structurally related proteins: this included the periplasmic dipeptide-binding proteins DppA from *Helicobacter pylori* (PDB 6PU3, sequence ID = 28.5%) (49) **(Figure 3D),** DppA from *E. coli* K-12 (PDB 1DPP, sequence ID= 30.7%) (50), the dipeptide transport protein from *Yersinia pestis* (PDB 5F1Q, sequence ID = 30 %) (51), and the glutathione-binding protein A (GbpA) from *Haemophilus parasuis* SH0165 (PDB 3M8U, sequence ID = 30%) (23) **(Supplementary Figure 5).** The binding sites of these proteins and their mode of binding to the ligands share similar characteristics in that they all have conserved residues at positions equivalent to Trp410 and Asp413, which interact directly with the shared NH_2_ group of their ligands **(Supplementary Figure 5).** However, notable differences in the *Me*Ami_SBP structure include an insertion near residue Met408 that appears to disrupt the beta strand observed in other published SBP_bac_5 structures and facilitate binding to l-amino acid amides only. The insertion positions the backbone carbonyl of Met408 so that it can interact with the NH_2_ group of l-serinamide and, along with several additional bulky residues in *Me*Ami_SBP, such as Trp435, Tyr32 and Trp34, also appears to decrease the overall size of the binding pocket compared to the other ligand-bound SBP_bac_5 proteins. This may prevent the binding of larger ligands such as dipeptides. The binding site structure thus justifies the divergence of the ligand binding partners of *Me*Ami_SBP compared to otherwise highly structural homologous proteins.

It is important to note that SBPs are dynamic proteins that often sample both closed states (in which the subdomains are close together) and open states (in which the subdomains are further apart) in solution, and that the structure of *Me*Ami_SBP bound to l-serinamide likely represents the closed conformation of the protein. Indeed, an AF2-predicted model of apo-*Me*Ami_SBP shows a more open conformation than what is observed in the crystal structure of the *Me*Ami_SBP in complex with l-serinamide (**Supplementary Figure 6**) and may represent a physiologically-relevant open, unliganded state. Considering that the l-serinamide and the other l-amino acid amide ligands form bridging interactions with residues from the two core domains of *Me*Ami_SBP, it is likely that ligand binding would stabilize the closed conformation of the protein that is observed in the crystal structure.

### *Me*Ami_SBP has a Ca^2+^ binding site

The crystal structure of *Me*Ami_SBP also revealed clear electron density near residues Asp45, Tyr207 and Asp49 that was consistent with a metal ion binding site (**Figure 4A, Supplementary Figure 7**). Coordination geometry (pentagonal bipyramidal), environmental B-factors and bond lengths were most consistent with this being a Ca^2+^ ion (**Supplementary Figure 8**). Ca^2+^ binding to His_6_-tagged *Me*Ami_SBP was subsequently confirmed using inductively coupled plasma mass spectrometry (ICP-MS) and isothermal titration calorimetry (ITC). Specifically, calcium was detected by ICP-MS at 17.8 parts per billion in a purified sample of 10 μM *Me*Ami_SBP (**Figure 4B**). While high concentrations of nickel and zinc were also detected in this sample of His_6_-tagged *Me*Ami_SBP, the presence of nickel and zinc could be attributed to these metal ions binding the His_6_-tag rather than *Me*Ami_SBP: removal of the His_6_-tag abolished nickel, zinc and copper binding **(Supplementary Figure 9**). Further, ITC titrations showed that Ca^2+^ binds His_6_-cleaved *Me*Ami_SBP with high affinity (*K*_D_ 2.80 μM) **(Figure 4C**), which aligns with the metal coordination properties of another calcium-coordinating SBP in SBP_bac_5 (53) and similar (low μM *K*_D_) metal-binding affinity identified for the primary metal cofactor of different class members in the broader superfamily (54,55). While Ca^2+^ increased ligand-induced thermal shifts for L-alaninamide, glycinamide and L-tyrosinamide, it slightly decreased the thermostability of apo-*Me*Ami_SBP (**Figure 4D**). This may be attributed to altered structural dynamics that decrease thermostability, which is consistent with the increase in *Me*Ami_SBP entropy conferred by Ca^2+^ binding observed using ITC. These data suggest a role for Ca^2+^ as a structural metal ion in ligand-bound *Me*Ami_SBP.

**Figure 4.**
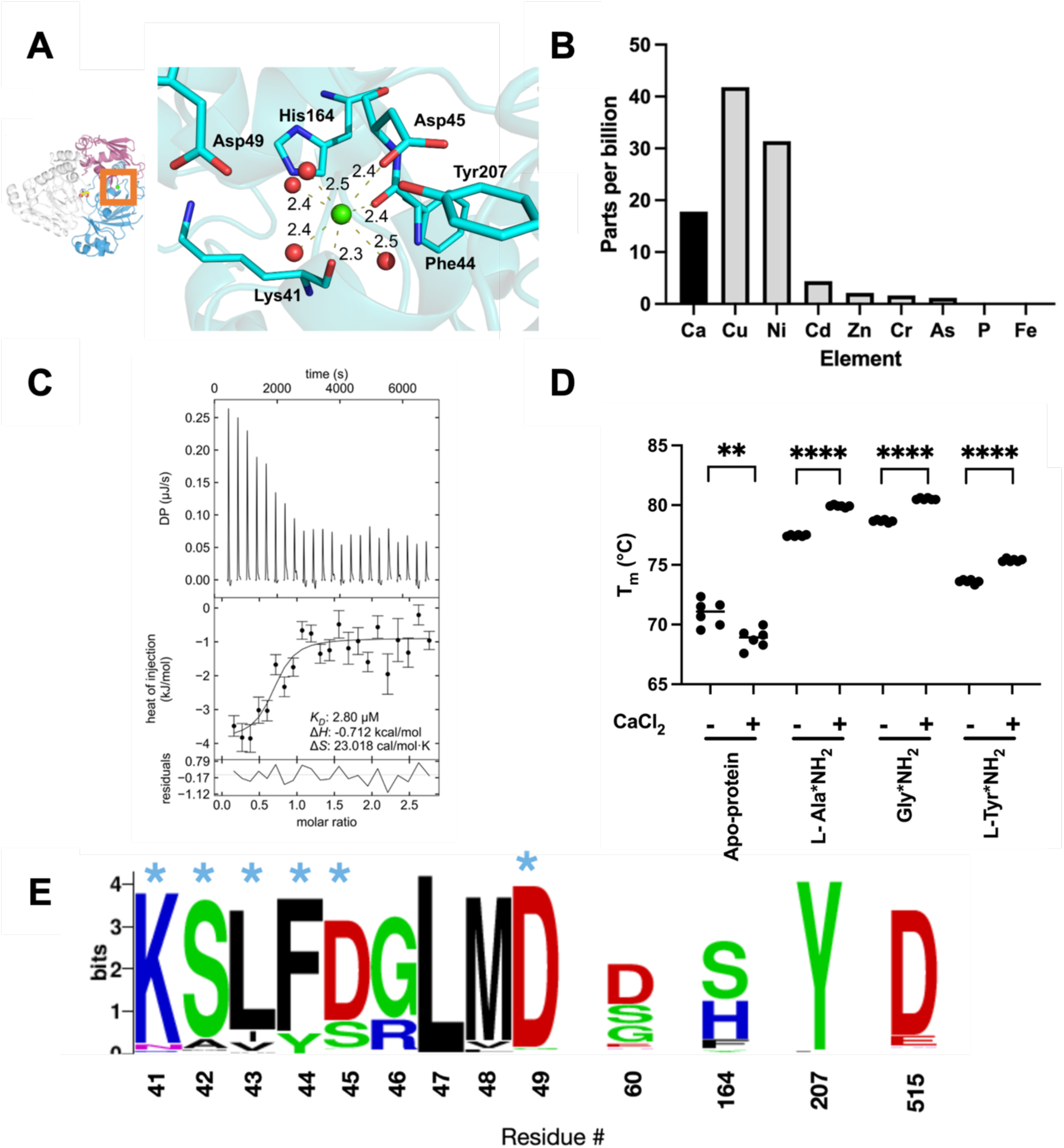
A conserved Ca^2+^ binding site in *Me*Ami_SBP. **A.** Ca^2+^-binding site coordination geometry and distances (in Å) in the *Me*Ami_SBP crystal structure. **B.** ICP-MS analysis of the inorganic elemental content of *Me*Ami_SBP directly after SEC purification. **C.** ITC thermogram of His_6_-cleaved *Me*Ami_SBP titrated with CaCl_2_ after removal of copurified Ca^2+^ ions by dialysis using EDTA. **D.** Protein melting temperatures measured using DSF for *Me*Ami_SBP in the presence of several amino acid amides ± 1 mM CaCl_2_. **** refers to p < 0.0001 and ** refers to p < 0.01 from unpaired parametric t-tests. **E.** Sequence WebLogo (52) showing conservation of residues within the Ca^2+^ coordination site in all homologs found in the same sequence cluster as *Me*Ami_SBP. Residues whose sidechains directly interact with the Ca^2+^ or the coordinating water molecules in *Me*Ami_SBP are highlighted with an asterisk.

In *Me*Ami_SBP, Ca^2+^ is coordinated by the side chain of Asp45, backbone carbonyl of Lys41, backbone carbonyl of Phe44, and four crystallographic waters that are held in place by the backbones and side chains of the surrounding residues (Ser42, Tyr207, Asp515, Asp49, Asp60 and His164). These residues are highly conserved across the other 609 SBPs found in the cluster with *Me*Ami_SBP (**Figure 4E**), suggesting that homologous SBPs probably bind Ca^2+^ at this position as well.

## DISCUSSION

The SBP_bac_5 family comprises periplasmic SBPs that are associated with ABC transport systems and that bind a range of substrates, including peptides, heme, nickel, opines, and other molecules (20–25). Our sequence similarity network analysis of this family highlighted that most of the sequence and function space of this family remains uncharacterized (**Figure 1**). In this work, we focused on a set of 610 uncharacterized proteins within the SBP_bac_5 family that are all found in conserved operons comprising genes encoding ABC transport systems and a putative amidase from the FmdA_AmdA family. Considering amides represent a valuable source of carbon and nitrogen for bacteria, and our understanding of import systems specific for such compounds is poor, we aimed to determine the ligand specificity of these homologous SBPs to provide insight into these amidase-associated uptake systems.

We found that although *Me*Ami_SBP, the most consensus-like representative from this subgroup, is annotated as a peptide-binding protein, it showed no ligand-induced thermal shifts in the presence of any of the di- or tri-peptides screened (**Figure 2**). We experimentally identified that *Me*Ami_SBP instead binds the l-amino acid amides l-serinamide, glycinamide, L-tyrosinamide and L-alaninamide with high (nM-μM *K*_D_) affinity (**Figure 2**). The structure of *Me*Ami_SBP highlights the structural determinants for specificity for the amidated amino acids over the canonical l-amino acids and other amide-containing molecules. Distinct specificity for amino acid amides is conferred through a combination of (i) a salt bridge between the N-terminal amine and Asp413, (ii) hydrogen bonds between the C-terminal carbonyl and Trp34 and Ile411 and (iii) hydrogen bonds between the carboxamide group, Tyr32 and Met408 (**Figure 3**). This three-component interaction determines ligand specificity, and ligand affinity is further tuned across two orders of magnitude based on side chain interaction compatibility with a hydrophobic binding site pocket, best highlighted by the differences in affinity between glycinamide and L-tyrosinamide. *Me*Ami_SBP likely diverges from its computational annotation as a peptide-binding protein due to a loop insertion in the binding site that reduces the volume of the pocket and occludes larger molecules that bind to homologs such as *H. pylori* DppA (49) **(Figure 3)**. However, it is important to note that *Me*Ami_SBP may also bind to other compounds not tested in this study that contain l-amino acid amide-like motifs. Most significantly, this work presents the first structural and experimental evidence for periplasmic SBP family proteins binding to l-amino acid amides. The conservation of binding site residues across the *Me*Ami_SBP cluster in the SSN indicates homologous binding properties exist in other bacteria.

The structure of *Me*Ami_SBP also revealed a Ca^2+^ binding site within its main small lobe, which was subsequently confirmed by both ITC and ICP-MS **(Figure 4)**. While the structural role of calcium is evident, its physiological significance in these proteins remains uncertain. Most significantly, this work presents the first evidence for SBP_bac_5 proteins implicated in calcium binding through multiple experimental methods, although calcium binding has been identified in other classes of the periplasmic SBP superfamily previously (56–58). The evidence for calcium binding having a structural role in *Me*Ami_SBP is compelling because the ion identified in the crystal structure was adventitiously bound during expression and proven to interact with *Me*Ami_SBP using ICP-MS and ITC. The conservation of coordination site residues across the *Me*Ami_SBP cluster in the SSN indicates calcium binding is a likely conserved property in the homologous proteins from other bacterial species.

The prevalence of Gram-negative nitrogen-fixing soil bacteria within this group, some of which are known to thrive on short chain amines (59), suggests an evolutionary advantage for soil bacteria capable of catalyzing amidase reactions on l-amino acid amides. This would metabolize them to the respective l-amino acid and ammonia, facilitating access to amino acid utilization pathways such as the citric acid cycle (60) and ammonia assimilation pathways such as the glutamine synthetase-glutamate synthase pathway (61), respectively. Characterizing the substrate specificity of the amidases found within these operons would help provide further insight into amide metabolism in these bacteria. This might also identify enzymes capable of producing enantiomerically pure l-amino acids from racemic amino acid amides.

The conserved operon structure and the presence of ANTAR protein and an additional cytosolic PBP in the operon also hints that the expression of these genes may be regulated via an ANTAR-dependent mechanism, like that of the amidase operon in *Pseudomonas aeruginosa* (35,36,41). Sequence and structural analyses of *Me*Ami_ANTAR and *Me*Ami_cPBP shows similarity to the *P. aeruginosa* AmiC/AmiR complex, but further research is needed to identify whether it responds to l-amino acid amides and to confirm the ANTAR target sequences, as canonical dual hexaloop motif sequences (62) could not be identified in this work. Future studies should determine if the system is induced by l-amino acid amides or other amide-containing compounds. This will expand our understanding of ANTAR regulated ABC transporter operons.

Overall, these findings contribute to our understanding of ABC transporter-associated SBPs binding to amides and their regulation in bacteria. The most compelling aspect of this work is that we provide the first experimental evidence for a cluster of periplasmic SBPs that bind l-amino acid amides. This evidence, in conjunction with the genetic elements co-located with *Me*Ami_SBP and sequence homology to co-clustered proteins from the source SSN, expands our understanding of microbial interactions with l-amino acid amides and general bacterial nitrogen utilization.

## METHODS

Key chemicals, reagents, recipes, and software are listed in Supplementary Information (**Supplementary Table 8-10**).

### Sequence Similarity Network and Bioinformatics

A sequence similarity network (SSN) of Pfam PF00496 (“Bacterial extracellular solute-binding proteins, family 5 Middle”) was calculated using EFI-EST (28) with the UniRef50 database. Default input settings were used. After the initial calculation, networks were calculated at a range of alignment score thresholds, with no sequence length restrictions. SSNs were visualized using the YFiles Organic Layout in Cytoscape 3.9.1 (63). The stringency of the SSN was increased until all unique SWISS-PROT descriptions were sorted into separate clusters of the SSN; this occurred at an alignment score of 100. The resulting isofunctional SSN was submitted to the EFI-GNT server (46) for analysis of per-cluster genomic context. *Me*Ami_SBP was selected as a representative of the target cluster using HMMER (43) and a hidden Markov model calculated from the MAFFT (64) alignment of all sequences in the cluster of interest.

### Protein expression and purification

SignalP 6.0 (65) was used to identify the signal peptide cleavage site between residues 25 and 26 of *Me*Ami_SBP sequence (probability=0.97) (**Supplementary Figure 10**). The signaling peptide (MKKFLASTVAASALALMLGMTSARA) was removed from the sequence encoding *Me*Ami_SBP and a DNA sequence encoding an N-terminal His_6_ tag and TEV protease cleavage site (GSSHHHHHHSSGENLYFQG) was added (**Supplementary Table 11**). This was codon-optimized for expression in *E. coli*, synthesized and cloned between NdeI and XhoI sites in pET-29b(+) vectors by Twist Bioscience (**Supplementary Table 12**). The plasmid was transformed into chemically-competent *E. coli* BL21(DE3) (NEB) using the recommended heat-shock protocol and plated onto LB Miller agar supplemented with 50 mg mL^-1^ kanamycin before being incubated overnight at 37 °C.

A single colony was picked and used to inoculate a seed culture in 10 mL LB media supplemented with 50 mg mL^-1^ kanamycin. The seed culture was incubated at 37°C for 8-12 hours with shaking at 200 rpm and then diluted 1:100 to inoculate a 1 L culture of simple autoinduction media (5 g L^-1^ Yeast Extract, 20 g L^-1^ Tryptone, 5 g L^-1^ NaCl, 6 g L^-1^ Na_2_HPO_4_.7H_2_O, 3 g L^-1^ KH_2_PO_4_, 0.19 g L^-1^ MgCl_2_, 6 g L^-1^ glycerol, 0.5 g L^-1^ glucose, 2 g L^-1^ D-lactose) supplemented with 50 mg mL^-1^ kanamycin in Thomson Ultra Yield Flasks (66). This was incubated at 30°C for 24 hours with shaking at 250 rpm. Cell pellets were harvested by centrifugation at 4730 rcf for 15 min at 20°C. Cell pellets were stored at −20 °C until purification.

*Me*Ami_SBP was purified using immobilized metal affinity chromatography (IMAC) followed by size exclusion chromatography (SEC). Cell pellets were resuspended in 25 mL equilibration buffer (20 mM sodium phosphate, 500 mM NaCl, 20 mM imidazole, pH 7.4). Cells were lysed by sonication (2 x 5 min at 50% power), followed by addition of 0.5 μL Turbonuclease from *Serratia marcescens* (Sigma Aldrich) and centrifugation at 17400 rcf for 75 min at 4°C to remove cell debris. The supernatant was filtered through a 0.45 μM syringe filter prior to loading onto a 5 mL HisTrap HP (Cytiva) column pre-equilibrated with equilibration buffer at 2.5 mL min^-1^. Following loading, the column was washed with 10 column volumes of equilibration buffer, followed by elution of the target protein with five columns of IMAC elution buffer (500 mM sodium phosphate, 500 mM NaCl, 20 mM imidazole, pH 7.4). Eluate fractions were pooled and loaded onto a size exclusion column (HiLoad Superdex 26/600 200 prep grade, GE Healthcare) pre-equilibrated with Tris buffer (50 mM Tris, 300 mM NaCl, pH 7.4) or HEPES buffer (50 mM HEPES, 300 mM NaCl, pH 7.4), depending on downstream use. SDS-PAGE (SurePAGE™, Bis-Tris, 10×8, 4-12%, GenScript) was used to assess protein purity, following which protein concentration was determined using a NanoDrop One^c^ Microvolume UV-Vis Spectrophotometer (ThermoFisher Scientific) using the predicted molar extinction coefficient from ProtParam (91,790 M⁻¹ cm⁻¹).

### Differential scanning fluorimetry (DSF)

For screening, Biolog Microbial Phenotyping Microarray Plates PM1-PM6 were resuspended in 125 μL MilliQ water and 5 μL of each well was aliquoted into MicroAmp Optical 96-Well Reaction Plates (ThermoFisher Scientific) to prepare screening plates. DSF experiments were performed in Tris buffer (50 mM Tris, 300 mM NaCl, pH 7.4) using a final protein concentration of 0.5 mg mL^-1^. A 1.33 × master mix was prepared containing *Me*Ami_SBP, Protein Thermal Shift buffer, Protein Thermal Shift dye and Tris buffer according to the Protein Thermal Shift™ Dye Kit (ThermoFisher Scientific) protocols. 15 μL of master mix was aliquoted into each well of the prepared screening plates to a final volume of 20 μL. Prepared plates were centrifuged at 2250 rcf for 1 min and sealed using MicroAmp Optical Adhesive Film (ThermoFisher Scientific). Melt experiments were performed using a QuantStudio Real-Time PCR thermocycler (ThermoFisher Scientific). Samples were heated from 25 °C to 95 °C at 0.05 °C s^-1^. Protein Thermal Shift Software (ThermoFisher Scientific) was used for data analysis. The midpoint of the unfolding transition was obtained from the maximum of the first derivative of fluorescence.

For testing of additional ligands, each ligand was prepared at 100 mM in Tris buffer and 2 μL was aliquoted into MicroAmp Optical 96-Well Reaction Plates (ThermoFisher Scientific). Plates were then prepared using 18 μL of a 1.11 × master mix according to the Protein Thermal Shift™ Dye Kit (ThermoFisher Scientific).

For isothermal analysis, 2 μL of serially diluted ligands were aliquoted into MicroAmp Optical 96-Well Reaction Plates (ThermoFisher Scientific). Plates were then prepared using 18 μL of a 1.11 × master mix according to the Protein Thermal Shift™ Dye Kit (ThermoFisher Scientific) recommendations. Protein concentration was 8.4 μM in each well. Data analysis was performed using FoldAffinity (48). For each isothermal analysis, the signal D window range was 40-85 °C. A global model was used to fit fluorescence with a ΔCP of 0 kcal K^-1^ mol^-1^, and a one-site model was used for the isothermal fit. Results were exported and plotted in GraphPad Prism.

### TEV Protease (TEVp)-mediated cleavage of N-terminal His_6_ tag

The His_7_-TEV (L56V, S135G, S219V) variant of the Tobacco Etch Virus protease (TEVp) was expressed and purified as described elsewhere (67). TEVp-cleavage of IMAC-purified His_6_-*Me*Ami_SBP was performed overnight in TEVp-cleavage buffer (10 mM Tris, 50 mM NaCl, 0.5 mM EDTA, 1 mM DTT, pH 8.0) using a protease to substrate mass ratio of 1:10. The following day, the cleaved substrate was recovered from the reaction by using a 5 mL HisTrap HP (Cytiva) column pre-equilibrated with equilibration buffer to remove uncleaved substrate, the cleaved His_6_-tag, and the His_7_-tagged protease, followed by size exclusion chromatography as described previously.

### Isothermal titration calorimetry (ITC)

His_6_-cleaved *Me*Ami_SBP was dialyzed extensively prior to ITC to remove Ca^2+^ ions and adventitiously bound ligand obtained via copurification: protein was dialyzed for 24 hours against 5 L of 50 mM HEPES, 300 mM NaCl, 10 mM EDTA, pH 7.4, followed by four passes against 5 L of 5 mM HEPES, 300 mM NaCl, pH 7.4, followed by one pass against 5 L of ITC buffer (50 mM HEPES, 300 mM NaCl, pH 7.4, 0.02% TWEEN20). Protein was concentrated to 10 mL using an Amicon Ultra-15 10K MWCO Centrifugal Filter Unit (Merck Millipore) and loaded into SnakeSkin Dialysis Tubing, 10K MWCO, 35 mm dry I.D. (ThermoFisher Scientific).

All titrations were performed in a TA Instruments Benchtop Nano ITC (small volume). All stocks and dilutions used for ITC were prepared using exactly matched buffer from the final round of dialysis. Proteins were concentrated using an Amicon Ultra-15 10K MWCO Centrifugal Filter Unit (Merck Millipore). Protein and ligand samples were degassed prior to each ITC experiment. All titrations were performed as titrations of ligand into protein at 25°C with a stirring rate of 350 rpm. Initial and final baselines were generated over 120 s. The first injection of each titration was a 1 μL dummy injection, followed by 22 injections of 2 μL each, with an injection interval of 300 s. The background heat was estimated as the average heat associated with each injection in a control titration of ligand into buffer and subtracted from each titration. Results were analyzed in NITPIC (68) for baseline detection, blank subtraction, and integration of baseline-subtracted power. SEDPHAT (69) was used to fit integrated heats to the single binding site model (A + B ←→ AB, heteroassociation) by global fitting with the Simplex algorithm. Thermograms were plotted in GUSSI (70).

### AlphaFold2 Structural Models

An AlphaFold2 model of *Me*Ami_SBP was generated using the “AlphaFold2_mmseqs2” ColabFold Notebook (71), which uses AlphaFold2 (42) and MMSeqs2 (72). The sequence corresponding to the TEVp-cleaved *Me*Ami_SBP protein was used. Default settings were used.

Structural figures were prepared using PyMOL Molecular Graphics System, Version 2.0 Schrödinger, LLC.

### Protein Crystallization and Data Collection

The His_6_-cleaved *Me*Ami was purified through size exclusion chromatography, as described above, using 4× crystallization buffer (40mM Tris, 150mM NaCl, pH 7.4). The purified protein sample was concentrated to 40 mg mL^-1^ using Amicon Ultra-15 3K MWCO Centrifugal Filter Unit (Merck Millipore) and subsequently diluted 1:4 with MilliQ water (yielding 10 mg mL^-1^ of protein in 1x crystal buffer (10mM Tris, 37.5 mM NaCl, pH 7.4). Sparse-matrix crystallography trials with ShotGun SG1 (Molecular Dimensions) and Index (Hampton Research) screens were performed at 18°C using sitting-drops of 400 nl protein mixed with 400 nl reservoir solution (prepared using a Formulatrix NT8).

A single large, cubic crystal formed after 3 months in well H5 of the ShotGun SG1 screen (30% w/v PEG 4000). The crystal was mounted onto a loop and set on goniometer without cryoprotecting or flash cooling. Diffraction data were collected in-house using an Agilent SuperNova X-ray source and processed using CrysAlisPro (Oxford Diffraction /Agilent Technologies UK Ltd, Yarnton, England) in space group P 2_1_ 2_1_ 2_1_. Data were truncated using Aimless (73) until statistics were of suitable quality in the outer shell. The structures were solved by molecular replacement (Phaser MR, CCP4 (74)) using the two domains of the AF2-predicted model of *Me*Ami_SBP (domain 1= residues 1-265, 488-523; domain 2= residues 266-488) as the search models. AF2 model pLDDT values were converted to B-factors in Phenix prior to molecular replacement. Refinement was carried out in Phenix.refine (75). Multiple rounds of refinement and model building in Coot (76) were carried out until R-factors converged and the structure was modeled as best as possible to the electron density. MolProbity (77) was used for structure validation. In the final structure, residue number 1 corresponds to the first residue (glycine) in the TEVp-cleaved *Me*Ami_SBP sequence. The metal binding site was analyzed using the CheckMyMetal (CMM): Metal Binding Site Validation Server (78). Polder omit maps were generated using phenix.polder (79) with default settings.

### FoldSeek

Similar structures were searched for using FoldSeek (80). All available databases were searched with Foldseek in 3Di/AA mode.

### Computational Docking of Ligands into *Me*Ami_SBP

Schrödinger (Schrödinger Release 2022-1: Desmond Molecular Dynamics System, D. E. Shaw Research, New York, NY, 2021.) was used for computational docking. Briefly, the crystal structure of *Me*Ami_SBP was prepared using default settings in the protein preparation wizard (pH 7.4, including a minimization step) and the ligands were prepared using the LigPrep wizard and default settings (retaining stereochemistry). The ligands were then docked into the *Me*Ami_SBP using the induced fit protocol with default settings. The OPLS4 Force Field (81) was used.

### Inductively Coupled Plasma Mass Spectrometry (ICP-MS)

ICP-MS was performed by technical staff at the Australian National University Research School of Earth Sciences ICP-MS Research Facility. 5 mL of 10 μM *Me*Ami_SBP was prepared in MS buffer (50 mM HEPES, 300 mM NaCl, pH 7.4). Both sample and suspension blank were diluted 1:10 in triplicate using 2% HNO_3_ to acidify samples and precipitate protein component. One of each triplicate was spiked with 100 ppb of Agilent Intelliquant 68 multi-element standard No.1 for ICP (IQ1), 5% HNO_3_, 100 μg mL^-1^ and 100 ppb of Agilent Intelliquant 12 multi-element standard No.2 for ICP, 5% HNO3, 100 μg mL^-1^ (IQ2). Solutions were first run on an Agilent 5110 OES to determine elemental composition of solutions. Samples were then run on a ThermoFisher iCapRQ Q-ICP-MS, excluding Na because this was very high in the suspension liquid. Final concentrations were processed by suppressing the 100 ppb addition of IQ1 and IQ2 and subtracting components of the blank sample.

## DATA AVAILABILITY

The 1.5 Å P 2_1_ 2_1_ 2_1_ crystal structure of the *Me*Ami_SBP–l-serinamide complex was submitted to the Protein Data Bank (PDB ID 8UPI). Raw data and associated files are available at doi: 10.5281/zenodo.10070929. Any other data is available upon request.

## AUTHOR CONTRIBUTIONS

OBS performed protein expression, purification, and experiments. RLF performed crystallography experiments, collected X-ray diffraction data, and processed the data. MGR performed protein expression, purification and initial DSF experiments. OBS, JAK and CJJ wrote the manuscript. CJJ supervised the project and provided support throughout. All authors edited and contributed to the final manuscript. All authors have read and approved the final version of the manuscript.

## ACKNOWLEDGEMENTS

This research was conducted by the Australian Research Council Centre of Excellence in Synthetic Biology (project number CE200100029) and funded by the Australian Government. We thank the Australian Research Council Centre of Excellence in Synthetic Biology for seed funding that supported this project. We thank Brett Knowles and Robin Grunn at the Australian National University Research School of Earth Sciences ICP-MS Research Facility for performing the ICP-MS experiments and for helpful discussions. We thank Dr Michael Gardiner for assistance with X-ray crystallography.

