## Supplementary Information for "Identification and characterization of a bacterial periplasmic solute binding protein that binds L-amino acid amides"

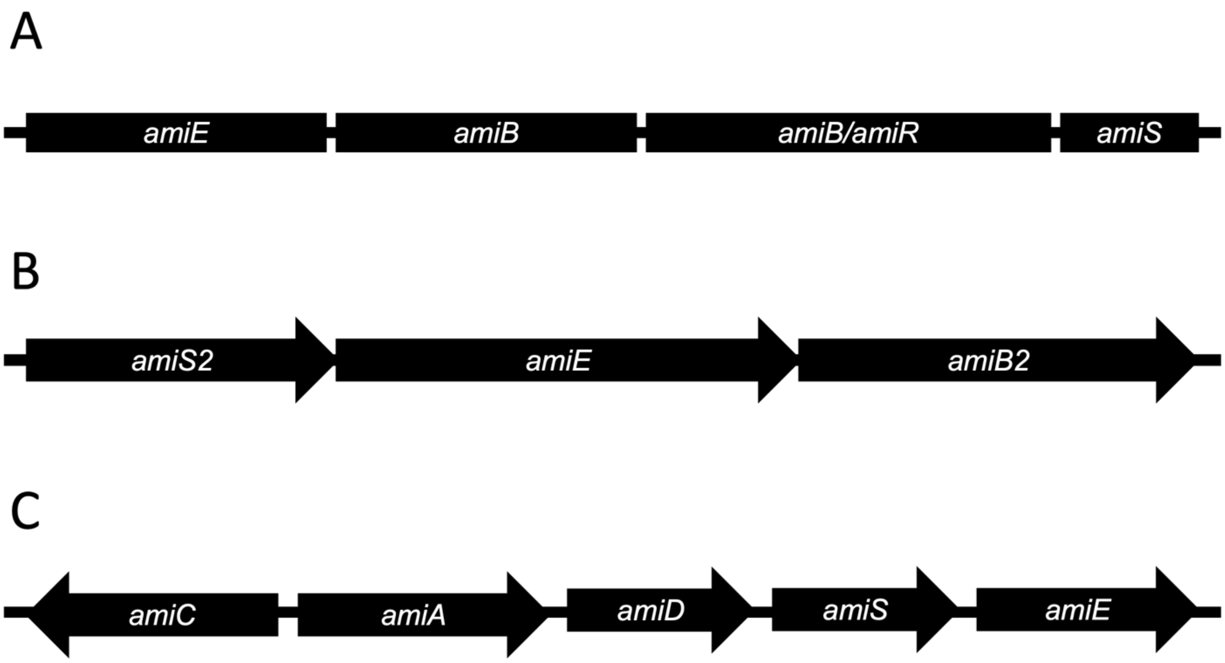


**Supplementary Figure 1. Amidase operon structure of characterised amide import systems. A.** *Pseudomonas aeruginosa* amidase operon (1,2), **B.** *Rhodococcus* sp. R312 amidase operon (3) and **C.** *Mycobacterium smegmatis* acetamidase operon (4–6). Presence, absence, and direction of arrows for each open reading frame is identical to operon structure reported in the referenced publications.

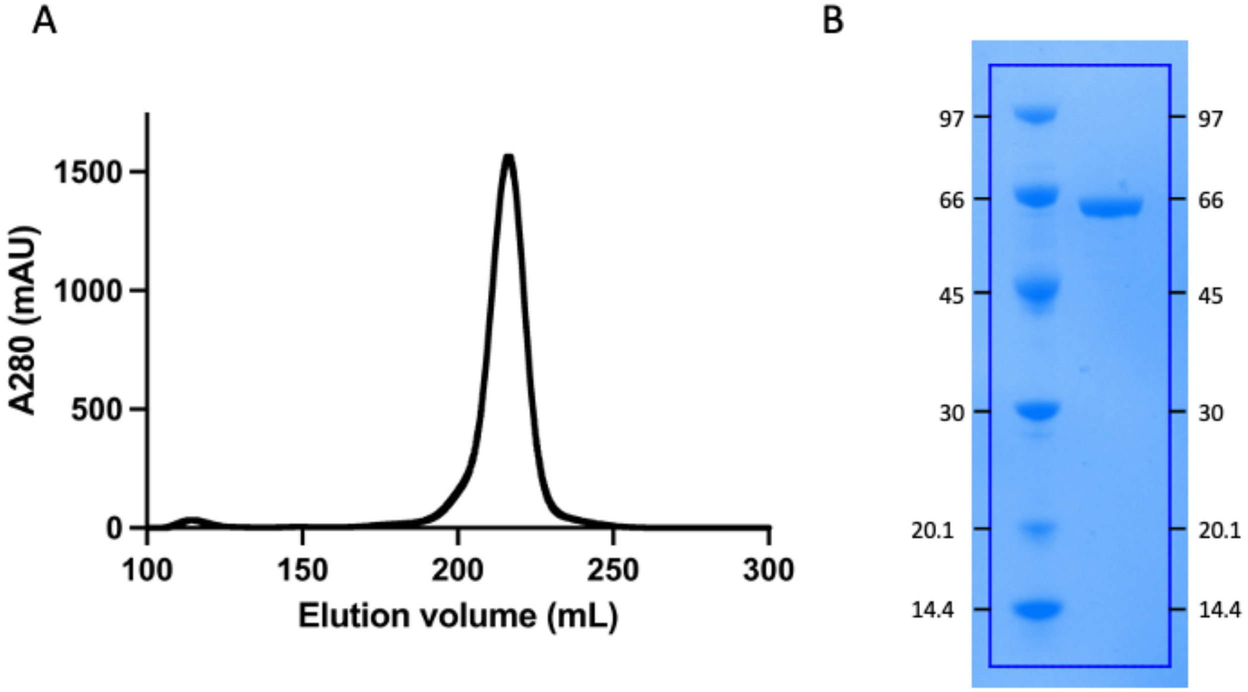


**Supplementary Figure 2.** (A) His$_6$-tagged *Me*Ami_SBP elutes from a HiLoad 26/600 Superdex 200 pg column consistent with it being monomeric in solution. (theoretical: 59.4 kDa (for His$_6$-tagged protein), calculated from standard curve: <60 kDa). (B) SDS-PAGE gel loaded with 1 µg His$_6$-tagged *Me*Ami_SBP showing size consistent with molecular weight relative to Amersham Low Molecular Weight Calibration Kit for SDS Electrophoresis.

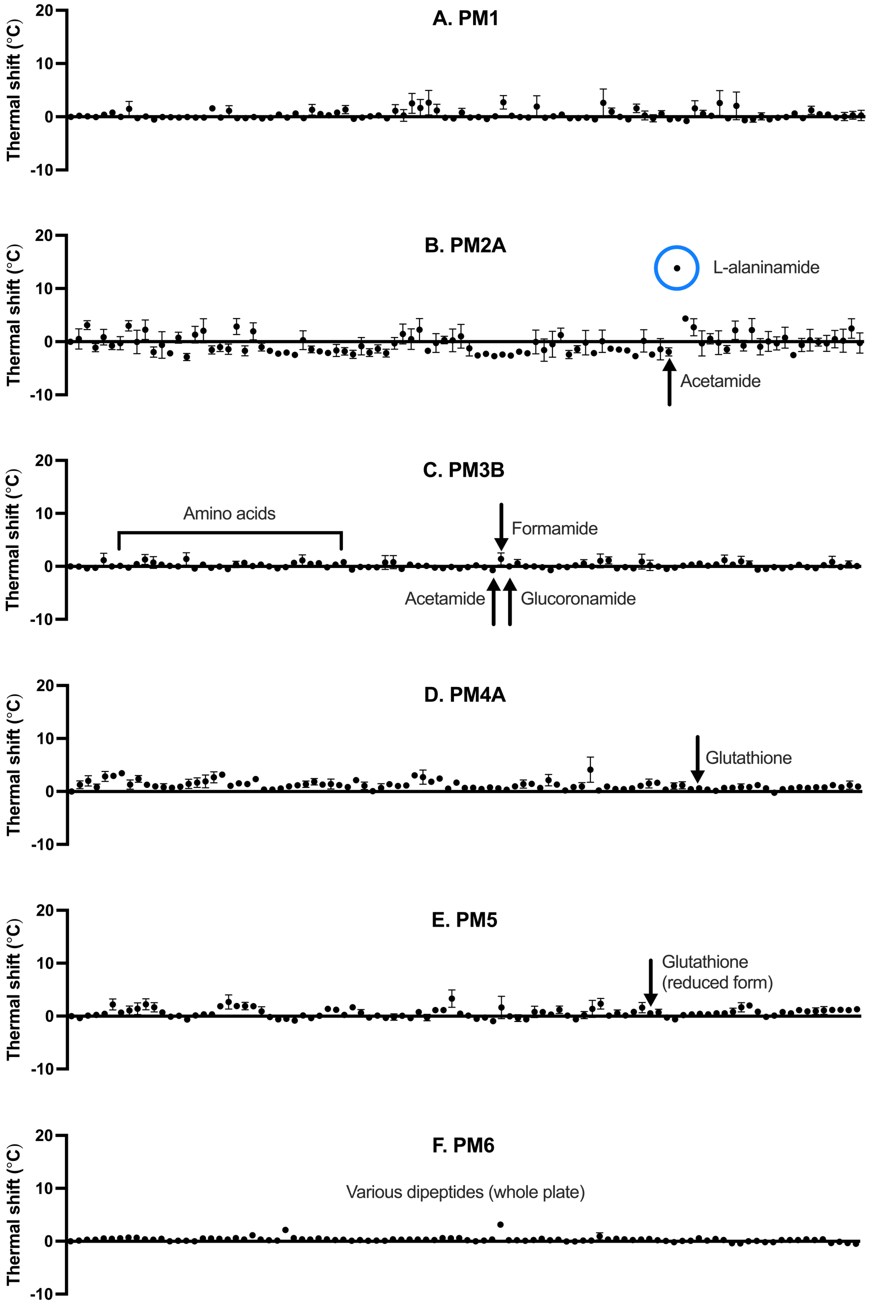


**Supplementary Figure 3.** Differential scanning fluorimetry ligand screen of *Me*Ami_SBP vs Biolog Microbial Phenotype MicroArray plates PM1-6 (A-F) (7), showing ligand-induced thermal shift compared with a no-ligand control. Each point represents a different ligand; ligand names hidden for clarity, with points representing L-alaninamide, L-alanine, and other key compounds highlighted. Error bars are standard error of the mean.

**Supplementary Table 1.** Biolog PM1 plate map.

| | | | | | | | | | | | |
|---|---|---|---|---|---|---|---|---|---|---|---|
| A1 Negative Control | A2 L-Arabinose | A3 N-Acetyl-D-Glucosamine | A4 D-Saccharic Acid | A5 Succinic Acid | A6 D-Galactose | A7 L-Aspartic Acid | A8 L-Proline | A9 D-Alanine | A10 D-Trehalose | A11 D-Mannose | A12 Dulcitol |
| B1 D-Serine | B2 D-Sorbitol | B3 Glycerol | B4 L-Fucose | B5 D-Glucuronic Acid | B6 D-Gluconic Acid | B7 D,L-α-Glycerol-Phosphate | B8 D-Xylose | B9 L-Lactic Acid | B10 Formic Acid | B11 D-Mannitol | B12 L-Glutamic Acid |
| C1 D-Glucose-6-Phosphate | C2 D-Galactonic Acid-γ-Lactone | C3 D,L-Malic Acid | C4 D-Ribose | C5 Tween 20 | C6 L-Rhamnose | C7 D-Fructose | C8 Acetic Acid | C9 α-D-Glucose | C10 Maltose | C11 D-Melibiose | C12 Thymidine |
| D-1 L-Asparagine | D2 D-Aspartic Acid | D3 D-Glucosaminic Acid | D4 1,2-Propanediol | D5 Tween 40 | D6 α-Keto-Glutaric Acid | D7 α-Keto-Butyric Acid | D8 α-Methyl-D-Galactoside | D9 α-D-Lactose | D10 Lactulose | D11 Sucrose | D12 Uridine |
| E1 L-Glutamine | E2 m-Tartaric Acid | E3 D-Glucose-1-Phosphate | E4 D-Fructose-6-Phosphate | E5 Tween 80 | E6 α-Hydroxy Glutaric Acid-γ-Lactone | E7 α-Hydroxy Butyric Acid | E8 β-Methyl-D-Glucoside | E9 Adonitol | E10 Maltotriose | E11 2-Deoxy Adenosine | E12 Adenosine |
| F1 Glycyl-L-Aspartic Acid | F2 Citric Acid | F3 m-Inositol | F4 D-Threonine | F5 Fumaric Acid | F6 Bromo Succinic Acid | F7 Propionic Acid | F8 Mucic Acid | F9 Glycolic Acid | F10 Glyoxylic Acid | F11 D-Cellobiose | F12 Inosine |
| G1 Glycyl-L-Glutamic Acid | G2 Tricarballylic Acid | G3 L-Serine | G4 L-Threonine | G5 L-Alanine | G6 L-Alanyl-Glycine | G7 Acetoacetic Acid | G8 N-Acetyl-β-D-Mannosamine | G9 Mono Methyl Succinate | G10 Methyl Pyruvate | G11 D-Malic Acid | G12 L-Malic Acid |
| H1 Glycyl-L-Proline | H2 p-Hydroxy Phenyl Acetic Acid | H3 m-Hydroxy Phenyl Acetic Acid | H4 Tyramine | H5 D-Psicose | H6 L-Lyxose | H7 Glucuronamide | H8 Pyruvic Acid | H9 L-Galactonic Acid-γ-Lactone | H10 D-Galacturonic Acid | H11 Phenylethyl-amine | H12 2-Aminoethanol |

**Supplementary Table 2.** Biolog PM2A plate map.

| | | | | | | | | | | | |
|---|---|---|---|---|---|---|---|---|---|---|---|
| A1 Negative Control | A2 Chondroitin Sulfate C | A3 α-Cyclodextrin | A4 β-Cyclodextrin | A5 γ-Cyclodextrin | A6 Dextrin | A7 Gelatin | A8 Glycogen | A9 Inulin | A10 Laminarin | A11 Mannan | A12 Pectin |
| B1 N-Acetyl-D-Galactosamine | B2 N-Acetyl-Neuraminic Acid | B3 β-D-Allose | B4 Amygdalin | B5 D-Arabinose | B6 D-Arabitol | B7 L-Arabitol | B8 Arbutin | B9 2-Deoxy-D-Ribose | B10 i-Erythritol | B11 D-Fucose | B12 3-0-β-D-Galacto-pyranosyl-D-Arabinose |
| C1 Gentiobiose | C2 L-Glucose | C3 Lactitol | C4 D-Melezitose | C5 Maltitol | C6 a-Methyl-D-Glucoside | C7 β-Methyl-D-Galactoside | C8 3-Methyl Glucose | C9 β-Methyl-D-Glucuronic Acid | C10 α-Methyl-D-Mannoside | C11 β-Methyl-D-Xyloside | C12 Palatinose |
| D1 D-Raffinose | D2 Salicin | D3 Sedoheptulosan | D4 L-Sorbose | D5 Stachyose | D6 D-Tagatose | D7 Turanose | D8 Xylitol | D9 N-Acetyl-D-Glucosaminitol | D10 γ-Amino Butyric Acid | D11 δ-Amino Valeric Acid | D12 Butyric Acid |
| E1 Capric Acid | E2 Caproic Acid | E3 Citraconic Acid | E4 Citramalic Acid | E5 D-Glucosamine | E6 2-Hydroxy Benzoic Acid | E7 4-Hydroxy Benzoic Acid | E8 β-Hydroxy Butyric Acid | E9 γ-Hydroxy Butyric Acid | E10 a-Keto-Valeric Acid | E11 Itaconic Acid | E12 5-Keto-D-Gluconic Acid |
| F1 D-Lactic Acid Methyl Ester | F2 Malonic Acid | F3 Melibionic Acid | F4 Oxalic Acid | F5 Oxalomalic Acid | F6 Quinic Acid | F7 D-Ribono-1,4-Lactone | F8 Sebacic Acid | F9 Sorbic Acid | F10 Succinamic Acid | F11 D-Tartaric Acid | F12 L-Tartaric Acid |
| G1 Acetamide | G2 L-Alaninamide | G3 N-Acetyl-L-Glutamic Acid | G4 L-Arginine | G5 Glycine | G6 L-Histidine | G7 L-Homoserine | G8 Hydroxy-L-Proline | G9 L-Isoleucine | G10 L-Leucine | G11 L-Lysine | G12 L-Methionine |
| H1 L-Ornithine | H2 L-Phenylalanine | H3 L-Pyroglutamic Acid | H4 L-Valine | H5 D,L-Carnitine | H6 Sec-Butylamine | H7 D.L-Octopamine | H8 Putrescine | H9 Dihydroxy Acetone | H10 2,3-Butanediol | H11 2,3-Butanedione | H12 3-Hydroxy 2-Butanone |

**Supplementary Table 3.** Biolog PM3B plate map.

| | | | | | | | | | | | |
|---|---|---|---|---|---|---|---|---|---|---|---|
| A1 Negative Control | A2 Ammonia | A3 Nitrite | A4 Nitrate | A5 Urea | A6 Biuret | A7 L-Alanine | A8 L-Arginine | A9 L-Asparagine | A10 L-Aspartic Acid | A11 L-Cysteine | A12 L-Glutamic Acid |
| B1 L-Glutamine | B2 Glycine | B3 L-Histidine | B4 L-Isoleucine | B5 L-Leucine | B6 L-Lysine | B7 L-Methionine | B8 L-Phenylalanine | B9 L-Proline | B10 L-Serine | B11 L-Threonine | B12 L-Tryptophan |
| C1 L-Tyrosine | C2 L-Valine | C3 D-Alanine | C4 D-Asparagine | C5 D-Aspartic Acid | C6 D-Glutamic Acid | C7 D-Lysine | C8 D-Serine | C9 D-Valine | C10 L-Citrulline | C11 L-Homoserine | C12 L-Ornithine |
| D-1 N-Acetyl-L-Glutamic Acid | D2 N-Phthaloyl-L-Glutamic Acid | D3 L-Pyroglutamic Acid | D4 Hydroxylamine | D5 Methylamine | D6 N-Amylamine | D7 N-Butylamine | D8 Ethylamine | D9 Ethanolamine | D10 Ethylenediamine | D11 Putrescine | D12 Agmatine |
| E1 Histamine | E2 β-Phenylethyl-amine | E3 Tyramine | E4 Acetamide | E5 Formamide | E6 Glucuronamide | E7 D,L-Lactamide | E8 D-Glucosamine | E9 D-Galactosamine | E10 D-Mannosamine | E11 N-Acetyl-D-Glucosamine | E12 N-Acetyl-D-Galactosamine |
| F1 N-Acetyl-D-Mannosamine | F2 Adenine | F3 Adenosine | F4 Cytidine | F5 Cytosine | F6 Guanine | F7 Guanosine | F8 Thymine | F9 Thymidine | F10 Uracil | F11 Uridine | F12 Inosine |
| G1 Xanthine | G2 Xanthosine | G3 Uric Acid | G4 Alloxan | G5 Allantoin | G6 Parabanic Acid | G7 D,L-α-Amino-N-Butyric Acid | G8 γ-Amino-N-Butyric Acid | G9 ε-Amino-N-Caproic Acid | G10 D,L-α-Amino-Caprylic Acid | G11 δ-Amino-N-Valeric Acid | G12 α-Amino-N-Valeric Acid |
| H1 Ala-Asp | H2 Ala-Gln | H3 Ala-Glu | H4 Ala-Gly | H5 Ala-His | H6 Ala-Leu | H7 Ala-Thr | H8 Gly-Asn | H9 Gly-Gln | H10 Gly-Glu | H11 Gly-Met | H12 Met-Ala |

**Supplementary Table 4.** Biolog PM4A plate map.

| | | | | | | | | | | | |
|---|---|---|---|---|---|---|---|---|---|---|---|
| A1 Negative Control | A2 Phosphate | A3 Pyrophosphate | A4 Trimeta-phosphate | A5 Tripoly-phosphate | A6 Triethyl Phosphate | A7 Hypophosphite | A8 Adenosine- 2'-monophosphate | A9 Adenosine- 3'-monophosphate | A10 Adenosine- 5'-monophosphate | A11 Adenosine- 2',3'-cyclic monophosphate | A12 Adenosine- 3',5'-cyclic monophosphate |
| B1 Thiophosphate | B2 Dithiophosphate | B3 D,L-α-Glycerol Phosphate | B4 β-Glycerol Phosphate | B5 Carbamyl Phosphate | B6 D-2-Phospho-Glyceric Acid | B7 D-3-Phospho-Glyceric Acid | B8 Guanosine- 2'-monophosphate | B9 Guanosine- 3'-monophosphate | B10 Guanosine- 5'-monophosphate | B11 Guanosine- 2',3'-cyclic monophosphate | B12 Guanosine- 3',5'-cyclic monophosphate |
| C1 Phosphoenol Pyruvate | C2 Phospho-Glycolic Acid | C3 D-Glucose-1-Phosphate | C4 D-Glucose-6-Phosphate | C5 2-Deoxy-D-Glucose 6-Phosphate | C6 D-Glucosamine-6-Phosphate | C7 6-Phospho-Gluconic Acid | C8 Cytidine- 2'-monophosphate | C9 Cytidine- 3'-monophosphate | C10 Cytidine- 5'-monophosphate | C11 Cytidine- 2',3'-cyclic monophosphate | C12 Cytidine- 3',5'-cyclic monophosphate |
| D1 D-Mannose-1-Phosphate | D2 D-Mannose-6-Phosphate | D3 Cysteamine-S-Phosphate | D4 Phospho-L-Arginine | D5 O-Phospho-D-Serine | D6 O-Phospho-L-Serine | D7 O-Phospho-L-Threonine | D8 Uridine- 2'-monophosphate | D9 Uridine- 3'-monophosphate | D10 Uridine- 5'-monophosphate | D11 Uridine- 2',3'-cyclic monophosphate | D12 Uridine- 3',5'-cyclic monophosphate |
| E1 O-Phospho-D-Tyrosine | E2 O-Phospho-L-Tyrosine | E3 Phosphocreatine | E4 Phosphoryl Choline | E5 O-Phosphoryl-Ethanolamine | E6 Phosphono Acetic Acid | E7 2-Aminoethyl Phosphonic Acid | E8 Methylene Diphosphonic Acid | E9 Thymidine- 3'-monophosphate | E10 Thymidine- 5'-monophosphate | E11 Inositol Hexaphosphate | E12 Thymidine 3',5'-cyclic monophosphate |
| F1 Negative Control | F2 Sulfate | F3 Thiosulfate | F4 Tetrathionate | F5 Thiophosphate | F6 Dithiophosphate | F7 L-Cysteine | F8 D-Cysteine | F9 L-Cysteinyl-Glycine | F10 L-Cysteic Acid | F11 Cysteamine | F12 L-Cysteine Sulfinic Acid |
| G1 N-Acetyl-L-Cysteine | G2 S-Methyl-L-Cysteine | G3 Cystathionine | G4 Lanthionine | G5 Glutathione | G6 D,L-Ethionine | G7 L-Methionine | G8 D-Methionine | G9 Glycyl-L-Methionine | G10 N-Acetyl-D,L-Methionine | G11 L- Methionine Sulfoxide | G12 L-Methionine Sulfone |
| H1 L-Djenkolic Acid | H2 Thiourea | H3 1-Thio-β-D-Glucose | H4 D,L-Lipoamide | H5 Taurocholic Acid | H6 Taurine | H7 Hypotaurine | H8 p-Amino Benzene Sulfonic Acid | H9 Butane Sulfonic Acid | H10 2-Hydroxyethane Sulfonic Acid | H11 Methane Sulfonic Acid | H12 Tetramethylene Sulfone |

**Supplementary Table 5.** Biolog PM5 plate map.

| | | | | | | | | | | | |
|---|---|---|---|---|---|---|---|---|---|---|---|
| A1<br>Negative Control | A2<br>Positive Control | A3<br>L-Alanine | A4<br>L-Arginine | A5<br>L-Asparagine | A6<br>L-Aspartic Acid | A7<br>L-Cysteine | A8<br>L-Glutamic Acid | A9<br>Adenosine-3',5'-cyclic monophosphate | A10<br>Adenine | A11<br>Adenosine | A12<br>2'-Deoxy Adenosine |
| B1<br>L-Glutamine | B2<br>Glycine | B3<br>L-Histidine | B4<br>L-Isoleucine | B5<br>L-Leucine | B6<br>L-Lysine | B7<br>L-Methionine | B8<br>L-Phenylalanine | B9<br>Guanosine-3',5'-cyclic monophosphate | B10<br>Guanine | B11<br>Guanosine | B12<br>2'-Deoxy Guanosine |
| C1<br>L-Proline | C2<br>L-Serine | C3<br>L-Threonine | C4<br>L-Tryptophan | C5<br>L-Tyrosine | C6<br>L-Valine | C7<br>L-isoleucine + L-Valine | C8<br>trans-4-Hydroxy L-Proline | C9<br>(5) 4-Amino-Imidazole-4(5)-Carboxamide | C10<br>Hypoxanthine | C11<br>Inosine | C12<br>2'-Deoxy Inosine |
| D1<br>L-Ornithine | D2<br>L-Citrulline | D3<br>Chorismic Acid | D4<br>(-)Shikimic Acid | D5<br>L-Homoserine Lactone | D6<br>D-Alanine | D7<br>D-Aspartic Acid | D8<br>D-Glutamic Acid | D9<br>D,L-α,ε-Diamino-pimelic Acid | D10<br>Cytosine | D11<br>Cytidine | D12<br>2'-Deoxy Cytidine |
| E1<br>Putrescine | E2<br>Spermidine | E3<br>Spermine | E4<br>Pyridoxine | E5<br>Pyridoxal | E6<br>Pyridoxamine | E7<br>β-Alanine | E8<br>D-Pantothenic Acid | E9<br>Orotic Acid | E10<br>Uracil | E11<br>Uridine | E12<br>2'-Deoxy Uridine |
| F1<br>Quinolinic Acid | F2<br>Nicotinic Acid | F3<br>Nicotinamide | F4<br>β-Nicotinamide Adenine Dinucleotide | F5<br>δ-Amino-Levulinic Acid | F6<br>Hematin | F7<br>Deferoxamine Mesylate | F8<br>D-(+)-Glucose | F9<br>N-Acetyl D-Glucosamine | F10<br>Thymine | F11<br>Glutathione (reduced form) | F12<br>Thymidine |
| G1<br>Oxaloacetic Acid | G2<br>D-Biotin | G3<br>Cyano-Cobalamine | G4<br>p-Amino-Benzoic Acid | G5<br>Folic Acid | G6<br>Inosine + Thiamine | G7<br>Thiamine | G8<br>Thiamine Pyrophosphate | G9<br>Riboflavin | G10<br>Pyrrolo-Quinoline Quinone | G11<br>Menadione | G12<br>m-Inositol |
| H1<br>Butyric Acid | H2<br>D,L-α-Hydroxy-Butyric Acid | H3<br>α-Keto-Butyric Acid | H4<br>Caprylic Acid | H5<br>D,L-α-Lipoic Acid (oxidized form) | H6<br>D,L-Mevalonic Acid | H7<br>D,L-Carnitine | H8<br>Choline | H9<br>Tween 20 | H10<br>Tween 40 | H11<br>Tween 60 | H12<br>Tween 80 |

**Supplementary Table 6.** Biolog PM6 plate map.

| | | | | | | | | | | | |
|---|---|---|---|---|---|---|---|---|---|---|---|
| A1<br>Negative Control | A2<br>Positive Control: L-Glutamine | A3<br>Ala-Ala | A4<br>Ala-Arg | A5<br>Ala-Asn | A6<br>Ala-Glu | A7<br>Ala-Gly | A8<br>Ala-His | A9<br>Ala-Leu | A10<br>Ala-Lys | A11<br>Ala-Phe | A12<br>Ala-Pro |
| B1<br>Ala-Ser | B2<br>Ala-Thr | B3<br>Ala-Trp | B4<br>Ala-Tyr | B5<br>Arg-Ala | B6<br>Arg-Arg | B7<br>Arg-Asp | B8<br>Arg-Gln | B9<br>Arg-Glu | B10<br>Arg-Ile | B11<br>Arg-Leu | B12<br>Arg-Lys |
| C1<br>Arg-Met | C2<br>Arg-Phe | C3<br>Arg-Ser | C4<br>Arg-Trp | C5<br>Arg-Tyr | C6<br>Arg-Val | C7<br>Asn-Glu | C8<br>Asn-Val | C9<br>Asp-Asp | C10<br>Asp-Glu | C11<br>Asp-Leu | C12<br>Asp-Lys |
| D1<br>Asp-Phe | D2<br>Asp-Trp | D3<br>Asp-Val | D4<br>Cys-Gly | D5<br>Gln-Gln | D6<br>Gln-Gly | D7<br>Glu-Asp | D8<br>Glu-Glu | D9<br>Glu-Gly | D10<br>Glu-Ser | D11<br>Glu-Trp | D12<br>Glu-Tyr |
| E1<br>Glu-Val | E2<br>Gly-Ala | E3<br>Gly-Arg | E4<br>Gly-Cys | E5<br>Gly-Gly | E6<br>Gly-His | E7<br>Gly-Leu | E8<br>Gly-Lys | E9<br>Gly-Met | E10<br>Gly-Phe | E11<br>Gly-Pro | E12<br>Gly-Ser |
| F1<br>Gly-Thr | F2<br>Gly-Trp | F3<br>Gly-Tyr | F4<br>Gly-Val | F5<br>His-Asp | F6<br>His-Gly | F7<br>His-Leu | F8<br>His-Lys | F9<br>His-Met | F10<br>His-Pro | F11<br>His-Ser | F12<br>His-Trp |
| G1<br>His-Tyr | G2<br>His-Val | G3<br>Ile-Ala | G4<br>Ile-Arg | G5<br>Ile-Gln | G6<br>Ile-Gly | G7<br>Ile-His | G8<br>Ile-Ile | G9<br>Ile-Met | G10<br>Ile-Phe | G11<br>Ile-Pro | G12<br>Ile-Ser |
| H1<br>Ile-Trp | H2<br>Ile-Tyr | H3<br>Ile-Val | H4<br>Leu-Ala | H5<br>Leu-Arg | H6<br>Leu-Asp | H7<br>Leu-Glu | H8<br>Leu-Gly | H9<br>Leu-Ile | H10<br>Leu-Leu | H11<br>Leu-Met | H12<br>Leu-Phe |

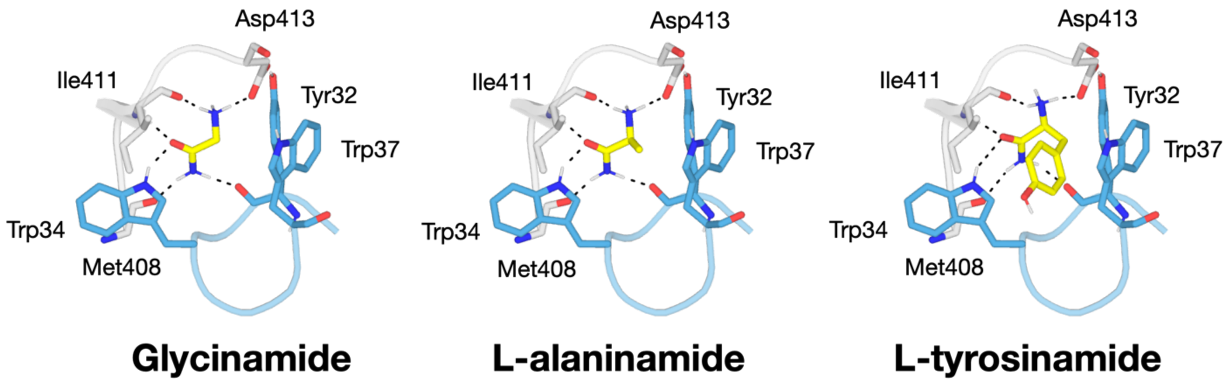


**Supplementary Figure 4.** Top-scoring poses from induced fit docking of glycinamide, L-alaninamide and L-tyrosinamide into *Me*Ami_SBP, showing identical backbone binding interactions and highlighting the ability of the *Me*Ami_SBP binding site to accommodate diverse ligand side chains. The two main subdomains of *Me*Ami_SBP are colored white (large domain) and blue (small domain), as in the figures presented in the main text.

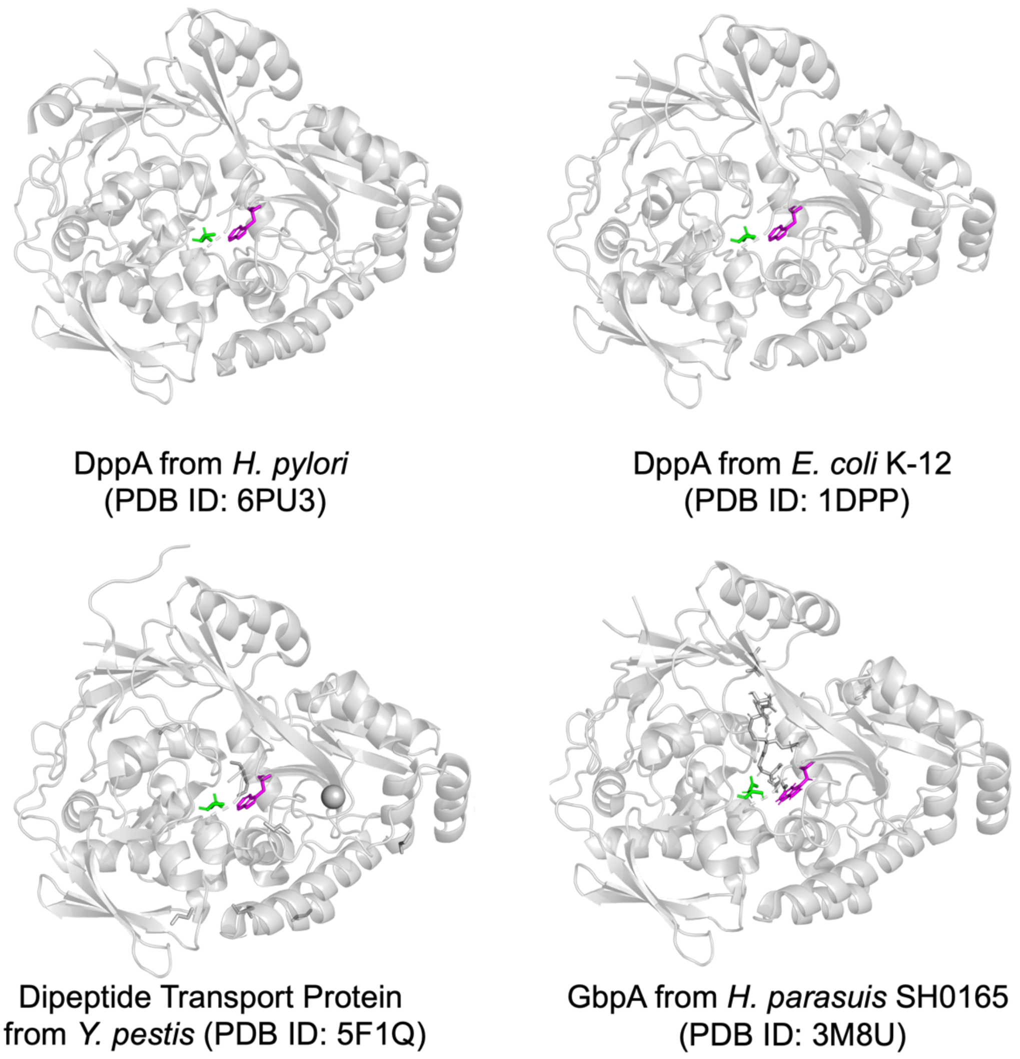


**Supplementary Figure 5**. Comparison of *Me*Ami_SBP binding site with other SBP_bac_5 structures (8–11) identifies identical residues equivalent to Trp410 (magenta) and Asp413 (green) in the binding site of each respective structure, indicating similar binding interactions with the $NH_3^+$ group of their respective ligands.

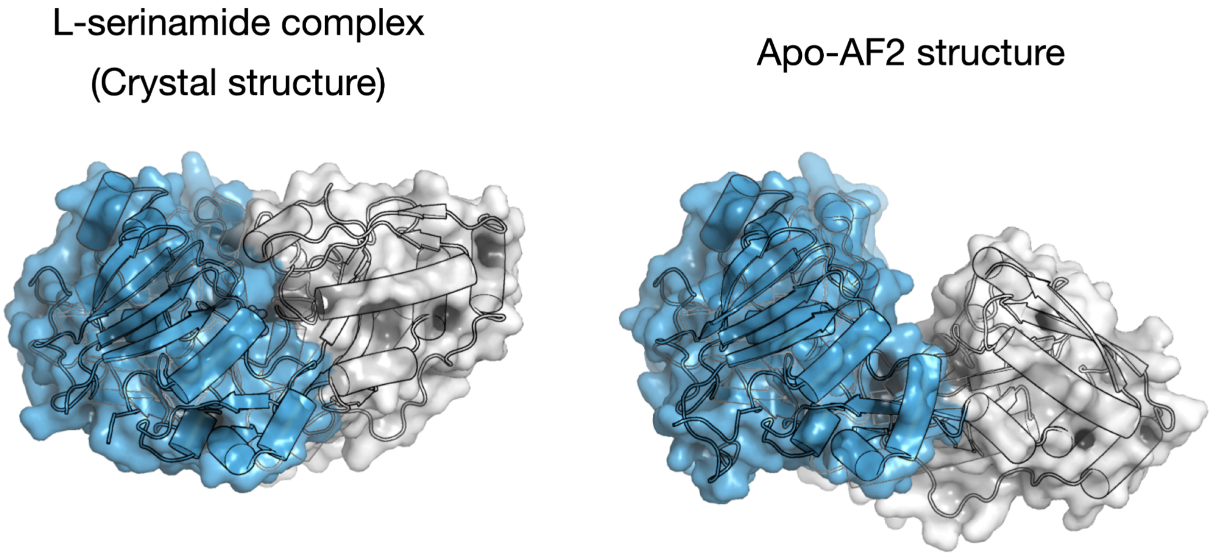


**Supplementary Figure 6.** Comparison of the crystal structure of *Me*Ami_SBP bound to L-serinamide (left) and the AlphaFold2-generated model of apo-*Me*Ami_SBP (12), highlighting possible open-closed rigid-body flexibility as is observed in other SBPs. The two key subdomains of *Me*Ami_SBP are colored blue and white for clarity.

**Supplementary Table 7. Crystallography statistics**

| PDB ID | 8UPI |
|---|---|
| **Data collection** | |
| Space group | P $2_1$ $2_1$ $2_1$ |
| **Cell dimensions** | |
| *a, b, c* (Å) | 56.25 58.92 139.35 |
| α, β, γ (°) | 90 90 90 |
| Resolution (Å) | 20.51 - 1.55 (1.58 - 1.55)* |
| $R_{merge}$ | 0.078 (0.503) |
| $R_{pim}$ | 0.041 (0.378) |
| I/σI | 7.8 (2.1) |
| $CC_{1/2}$ | 0.985 (0.445) |
| Completeness (%) | 99.2 (96.4) |
| Redundancy | 3.8 (2.9) |
| **Refinement** | |
| Resolution (Å) | 20.51 - 1.55 (1.57 - 1.55) |
| No. reflections | 66818 (6467) |
| $R_{work}/R_{free}$ | 0.1791 / 0.2285 (0.2825 / 0.3057) |
| **No. atoms** | |
| Protein | 4080 |
| Ligand/ion | 7 |
| Solvent | 859 |
| B-factors (overall) (Å$^2$) | 17.35 |
| Protein | 15.32 |
| Ligand/ion | 34.54 |
| Solvent | 26.78 |
| **R.M.S. deviations** | |
| Bond lengths (Å) | 0.005 |
| Bond angles (°) | 0.801 |

**Data in parentheses are for the highest resolution shell*

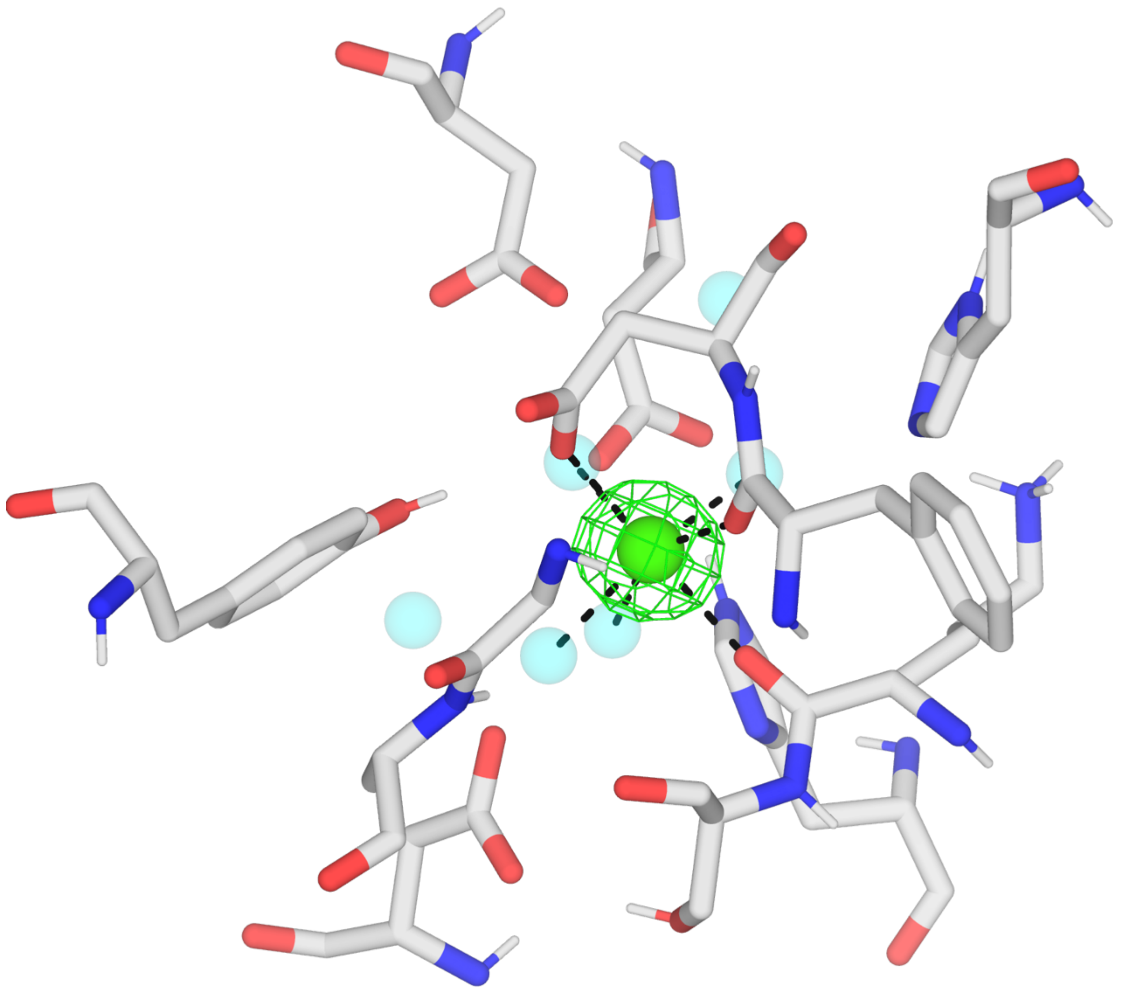

**Supplementary Figure 7. Polder omit map of metal binding site.** Polder omit map around the calcium ion (green sphere) generated using *phenix.polder* (13). Contoured at 3 sigma. 3 Å carve around ligand. Crystallographic water molecules are shown as cyan spheres.

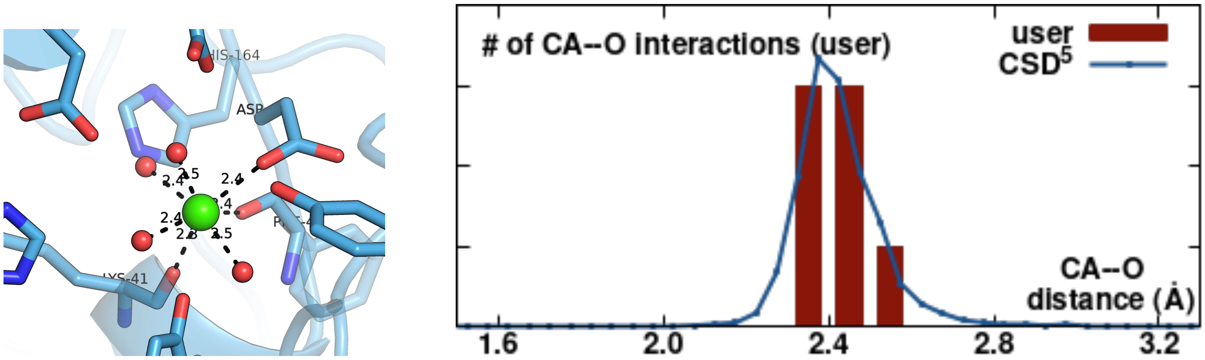


**Supplementary Figure 8. CheckMyMetal results.** CheckMyMetal (CMM): Metal Binding Site Validation Server (14) was used to assess the metal binding site. $Ca^{2+}$ was scored as the most probable metal ion based on coordination geometry (pentagonal bipyramidal, B-factor analysis and CA–O interaction distances (as shown by the plot on the right that compares the distances in the crystal structure (bars) with distances found in published structures).

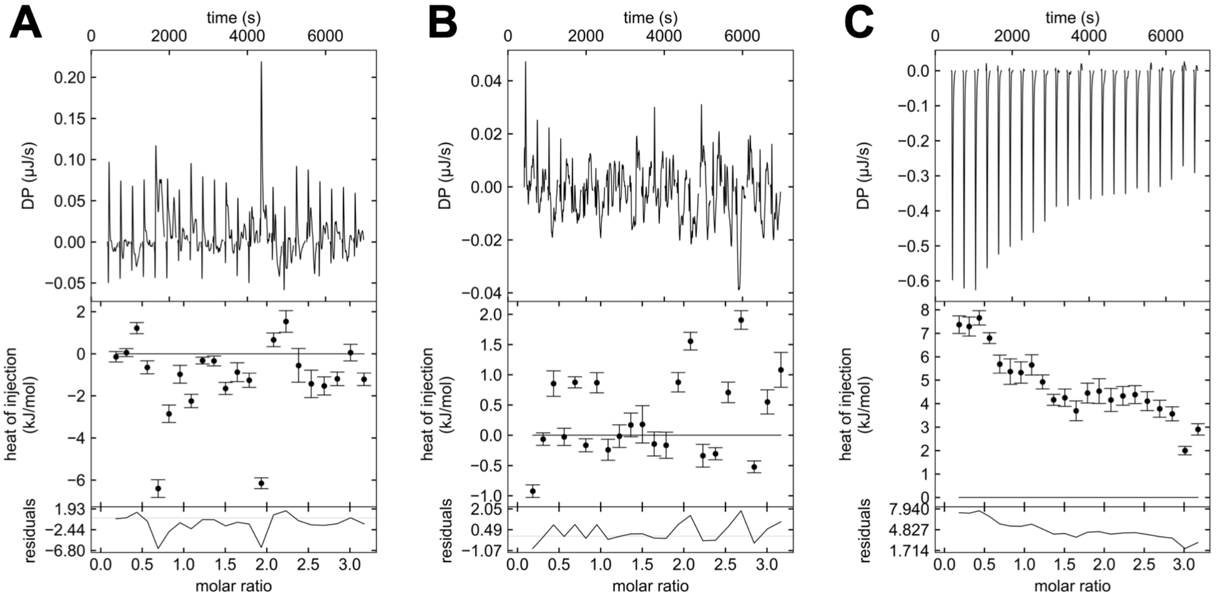


**Supplementary Figure 9. Isothermal titration calorimetry thermograms show that TEVp-mediated cleavage of the polyhistidine tag alleviates binding to copper, nickel and zinc ions. A.** Titration of *Me*Ami_SBP with $CuSO_4$ shows no heat of binding. **B.** Titration of *Me*Ami_SBP with $NiCl_2$ shows no heat of binding. **C.** Titration of *Me*Ami_SBP with $ZnSO_4$ shows only heat of dilution upon each injection.

**Supplementary Table 8. Key Materials Table**

| ITEM | Vendor | Catalog Number/link |
|---|---|---|
| Chemically-competent *Escherichia coli* BL21(DE3) | New England Biolabs | C2527I |
| Kanamycin monosulfate | AG Scientific | K-1022-25GM |
| Turbonuclease from *Serratia marcescens* | Sigma Aldrich | T4330 |
| 5 mL HisTrap HP | Cytiva | 17524802 |
| HiLoad Superdex 26/600 200 prep grade | GE Healthcare | GE28-9893-36 |
| SurePAGE, Bis-Tris, 10x8, 4-12% | GenScript | GB0746 |
| NanoDrop One$^{C}$ Microvolume UV-Vis Spectrophotometer | ThermoFisher Scientific | N/A |
| ThermoFisher Scientific Protein Thermal Shift Dye Kit | ThermoFisher Scientific | 4461146 |
| QuantStudio 3 Real-Time PCR thermocycler | ThermoFisher Scientific | N/A |
| Biolog Microbial Phenotype MicroArray (PM1, PM2A, PM3B, PM4A, PM5, PM6) | Biolog | 12111, 12112, 12121, 12131, 12141, 12181 |

| | | |
|---|---|---|
| MicroAmp Optical 96-Well Reaction Plate with Barcode | ThermoFisher Scientific | 4306737 |
| MicroAmp Optical Adhesive Film (ThermoFisher Scientific) | ThermoFisher Scientific | 4311971 |
| Protein Thermal Shift Software v1.3 (ThermoFisher Scientific) | ThermoFisher Scientific | 4466037 (v1.4) |
| Amicon Ultra-15 10K MWCO Centrifugal Filter Unit | Merck Millipore | UFC901024 |
| SnakeSkin Dialysis Tubing, 10K MWCO, 35 mm dry I.D. | ThermoFisher Scientific | 88245 |
| TA Instruments Benchtop Nano ITC | TA Instruments | N/A |
| Molecular Dimensions Shot Gun 1 Crystal Screen | Molecular Dimensions | N/A |
| Hampton Index HT Screen | Hampton Research | HR2-134 |
| Formulatrix NT8 crystallization robot | Formulatrix | N/A |

**Supplementary Table 9. Recipe Table**

| **Name** | **Contents** |
|---|---|
| LB Miller Media | 5 g $L^{-1}$ Yeast Extract, 10 g $L^{-1}$ Tryptone, 10 g $L^{-1}$ NaCl, |
| Simple autoinduction media (15) | 5 g $L^{-1}$ Yeast Extract, 20 g $L^{-1}$ Tryptone, 5 g $L^{-1}$ NaCl, 6 g $L^{-1}$ $Na_2HPO_4.7H_2O$, 3 g $L^{-1}$ $KH_2PO_4$, 0.19 g $L^{-1}$ $MgCl_2$, 6 g $L^{-1}$ glycerol, 0.5 g $L^{-1}$ glucose, 2 g $L^{-1}$ D-lactose |
| IMAC Equilibration Buffer | 20 mM sodium phosphate, 500mM NaCl, 20 mM imidazole, pH 7.4 |
| IMAC Elution Buffer | 500 mM sodium phosphate, 500 mM NaCl, 500 mM imidazole, pH 7.4 |
| Tris Buffer | 50 mM Tris, 300 mM NaCl, pH 7.4 |
| ITC Buffer | 50 mM HEPES, 300 mM NaCl, 0.02% TWEEN20, pH 7.4 |
| MS Buffer | 50 mM HEPES, 300 mM NaCl, pH 7.4 |
| Dialysis Buffer | 5 mM HEPES, 300 mM NaCl, pH 7.4 |
| Chelation buffer | 50 mM HEPES, 300 mM NaCl, 10 mM EDTA, pH 7.4 |
| 4 × crystal buffer | 40mM Tris, 150mM NaCl, pH 7.4 |

**Supplementary Table 10. Software Table**

| **Software** | **Version** |
| --- | --- |
| Protein Thermal Shift Software (ThermoFisher Scientific) | 1.3 |
| FoldAffinity (16) | N/A |
| NanoAnalyze | N/A |
| NITPIC (17) | 1.2.2 |
| SEDPHAT (18) | N/A |
| GUSSI (19) | 1.4.2 |
| Graphpad Prism | 9 |
| PyMOL | 2.5.2 |
| CCP4 – (including Aimless, Phaser MR) (20) | 7.1.018 |
| Phenix (21) | 1.20.1 |
| Schrödinger | Release 2022-1 |

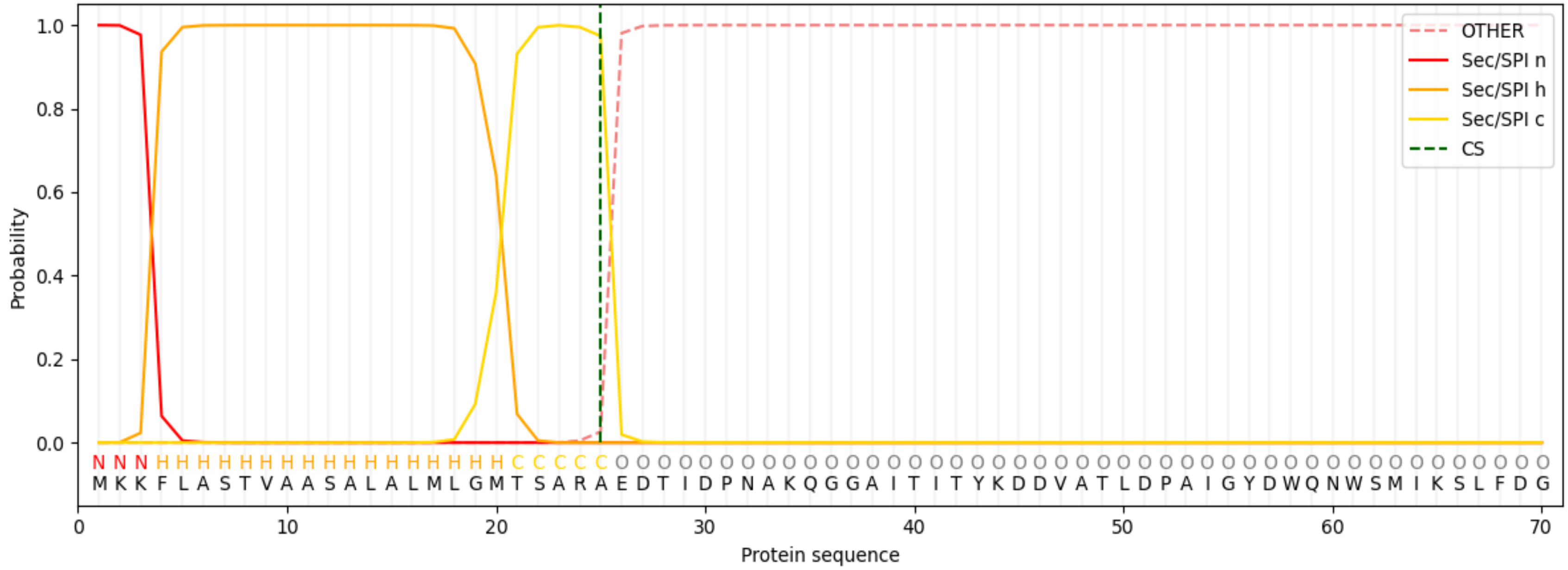


**Supplementary Figure 10.** Output from SignalP 6.0 (22), showing predicted Sec/SPI cleavage site between residues 25 and 26 of *Me*Ami_SBP (probability=0.97).

**Supplementary Table 11. Amino Acid Sequence of His$_6$-*Me*Ami_SBP.** Underlined + bold = TEVp Cut-site; RED = Sequence that is cleaved post TEVp-mediated cleavage; * = STOP CODON

| |
|---|
| MGSSHHHHHHSSG**<u>ENLYFQG</u>**EDTIDPNAKQGGAITITYKDDVATLDPAIGYDWQNWSM<br>IKSLFDGLMDYEPGTTNLKPDLAESYEISPDGKTFTFKLRHGVKFHNGREMTADDVKYS<br>LDRVTNPKTQSPGAGFFGSIKGYDDVAAGKATSLSGVTVVDPYTVKFELTRPDATFLHV<br>MAINFSHVVPKEEVEKYGADFGKHPVGTGAFKLAEWTLGQRIVFERNPDYWHKGLPHL<br>DKITFEIGQEPIVALLRLQKGEIDVPGDGIPPAKFQEVMADPEQKARVVEGGQLHTGYVT<br>MNTTMAPFDNVKVRQAVNMAINKARIIQIINGRAVPANQPLPPSMPGYDKEYKGYPYDV<br>AKAKALLAEAGHPDGFETQLFAMNTDPNPRIAQAIQQDLAAIGIKASIQSLAQANVIAAGG<br>DKAGAPMIWSGGMAWIADFPDPSNFYGPILGCAGAVPGGWNWSWYCNKDLDAKAAE<br>ADSVVDPAKGAERDKMWSAIYDKVMEDAPWAPVFNEQRFTMKSARMGGADNLYVDP<br>VHIPINYDNVYVKDVQ* |

**Supplementary Table 12. DNA Sequence encoding N-terminally $His_6$-tagged *Me*Ami_SBP.** This was inserted between NdeI and XhoI sites of pET-29b(+). The start codon (ATG) encoded by the vector is highlighted in red. Underlined region represents sequence encoding N-terminal $His_6$ and TEVp cut-site that was added to the sequence of *Me*Ami_SBP.

(ATG)GGAAGTTCGCATCATCATCATCATCACTCTTCAGGGGAAAATCTTTATTTTCAG
GGAGAAGACACCATAGACCCGAATGCGAAACAGGGTGGTGCAATTACAATTACATA
TAAAGATGATGTCGCGACCTTAGATCCGGCAATTGGCTATGATTGGCAGAACTGGT
CGATGATTAAGAGTCTTTTTGACGGACTCATGGATTACGAACCAGGGACTACAAATC
TGAAGCCCGATCTGGCTGAGAGCTATGAGATTTCGCCCGATGGGAAAACTTTCACG
TTCAAATTACGCCATGGTGTGAAGTTCCATAATGGGCGGGAAATGACAGCTGACGA
TGTAAAATATTCTCTGGACCGCGTAACCAATCCGAAAACGCAGTCTCCGGGAGCCG
GTTTCTTCGGCTCCATTAAGGGTTATGACGATGTGGCGGCAGGTAAAGCAACTTCTT
TGTCAGGGGTTACAGTAGTGGATCCCTATACTGTTAAATTCGAATTAACTCGTCCTG
ATGCAACGTTTTTACATGTAATGGCCATCAACTTTAGTCACGTTGTGCCGAAAGAAG
AGGTAGAGAAGTATGGAGCTGATTTTGGTAAACACCCGGTGGGTACGGGAGCCTTC
AAGCTGGCTGAGTGGACCCTGGGCCAGCGTATCGTTTTTGAACGGAATCCAGATTA
CTGGCATAAAGGCCTCCCGCACTTGGACAAAATTACGTTTGAAATCGGGCAGGAGC
CCATAGTAGCCCTGCTTCGACTCCAGAAAGGTGAAATTGATGTCCCTGGCGACGGT
ATCCCACCTGCTAAATTTCAAGAAGTAATGGCCGATCCAGAACAGAAAGCGCGCGT
GGTTGAGGGTGGTCAGCTGCATACCGGCTACGTTACCATGAACACCACAATGGCGC
CTTTTGATAATGTGAAAGTTAGACAGGCAGTCAACATGGCCATTAACAAGGCCCGCA
TAATACAAATCATAAACGGAAGAGCGGTTCCGGCAAACCAACCATTGCCGCCGTCA
ATGCCAGGATATGACAAAGAGTACAAAGGTTATCCGTATGATGTCGCTAAAGCGAAA
GCCCTGCTTGCTGAAGCAGGCCATCCTGATGGCTTTGAGACTCAATTATTTGCCATG
AATACCGACCCTAATCCTCGTATTGCTCAGGCTATCCAACAAGATCTGGCCGCGATA
GGCATTAAAGCTAGCATTCAAAGCCTGGCGCAGGCAAATGTCATCGCAGCGGGCG
GGGATAAAGCGGGTGCACCTATGATCTGGAGTGGGGGCATGGCATGGATCGCTGA
CTTTCCAGATCCATCTAACTTTTACGGACCTATTTTGGGCTGCGCGGGGGCGGTTC
CGGGAGGTTGGAATTGGTCCTGGTACTGTAATAAGGATCTGGATGCAAAAGCCGCT
GAAGCAGATAGCGTTGTCGATCCCGCTAAGGGTGCCGAACGTGACAAGATGTGGTC
AGCCATCTATGATAAAGTTATGGAAGATGCCCCGTGGGCACCCGTGTTTAATGAACA

GCGTTTCACGATGAAATCCGCCCGAATGGGCGGAGCGGACAACCTTTATGTGGATC
CAGTGCACATTCCGATTAATTATGACAATGTTTACGTGAAAGACGTCCAGTAA
